# 3D Hybrid Bioprinting for Complex Multi-Tissue Engineering

**DOI:** 10.1101/2025.11.06.682452

**Authors:** Hossein Vahid Alizadeh, Andrea S. Flores Pérez, Tomohiro Uno, Richard Sammy Muniz, Sun Hyung Kwon, Anjana Balachandar, Nicholas Riley, Chantal Aivan Le, Jiannan Li, Peng Zhao, Elaine Lui, Carolyn Kim, Seyedsina Moeinzadeh, Chi-Chun Pan, Nidhi Bhutani, Constance Chu, Sungwoo Kim, Yunzhi P. Yang

**Affiliations:** Department of Orthopaedic Surgery, Stanford University, Stanford, USA; Department of Bioengineering, Stanford University, Stanford, USA; School of Medicine, Stanford University, Stanford, USA; Department of Biology, Stanford University, Stanford, USA; Department of Mechanical Engineering, Stanford University, Stanford, USA; Department of Materials Science and Engineering, Stanford University, Stanford, USA

**Keywords:** bioprinting, multi-material biofabrication, tissue engineering, biomaterials, orthopedic grafts, musculoskeletal therapeutics

## Abstract

3D bioprinting has revolutionized tissue engineering, enabling intricate, physiologically relevant constructs unattainable with conventional techniques, yet it remains limited in integrating soft and rigid multifunctional components for complex multi-tissue applications. In this study, we introduce a 3D hybrid bioprinting approach implementing the Hybprinter platform, which integrates multiple 3D printing modules under optimized conditions for a continuous bioprinting process with multiple soft and hard biomaterials. This approach demonstrates robust biocompatibility and broad tissue engineering potential for modeling and therapeutic applications. The capacity to fabricate multi-hydrogel hybrid constructs is illustrated by representative examples highlighting vascularization, multifunctionality, mechanical robustness, and implant suturability. Notably, compared with commonly fabricated hydrogel-only constructs, the resulting hybrid constructs achieve over a 1000-fold increase in mechanical strength, and demonstrated enhanced osteogenic differentiation, underscoring their suitability for load-bearing musculoskeletal and orthopedic tissue engineering. Additionally, cell-laden hydrogel constructs demonstrated robust chondrogenic differentiation, highlighting the capacity for lineage-specific tissue development in vitro. Beyond these outcomes, the presented hybrid bioprinting approach integrates essential tissue engineering attributes that unites mechanical robustness and suturable capacity with multi-material integration, gradient property design, incorporation of bioactive agents, and support for multi-cell loading. This versatile platform advances complex tissue engineering and holds promise for patient specific, organ-on-demand applications.

## 1. Introduction

The field of tissue engineering focuses on developing tissues and organs to address two main clinical needs: creating more representative in vitro disease models and treating tissue defects and organ loss in patients^1,2^. Due to the global organ shortage crisis and aging population, there is an increasing demand for novel tissue engineering methods that enable organ-on-demand applications that can mimic native tissues both in structure and function while also reducing the need for animal studies. 3D bioprinting has emerged as a core technology for addressing these clinical needs through additive manufacturing of cellular constructs that can serve as an in vitro platform for applications in disease modeling^3^, drug screening and delivery^4,5^, cancer therapies^6,7^, food industry^8^, tissue engineering, and regenerative medicine^9^. In recent years, multiple approaches have been optimized to create more physiologically relevant tissue constructs, such as inkjet bioprinting, syringe-based microextrusion (SBM), molted material extrusion (MME), stereolithography in conjunction with digital light processing (DLP-SLA), and laser-assisted bioprinting. Each of these technologies offers its own advantages and limitations in engineering biological tissues. For instance, DLP-SLA and laser-assisted bioprinting are nozzle-free and provide higher resolution printing compared to inkjet and SBM methods – but inkjet and SBM are more cost-effective and user-friendly. MME offers enhanced load-bearing and surgical handling properties such as suturing capacity compared to all other bioprinting approaches – but cannot be loaded with cells nor mimic soft tissue composition like inkjet, SBM, and DLP-SLA bioprinting. Overall, each technique offers unique benefits to bioprinting through their methods and materials, but also face limitations in achieving the biological, mechanical, and geometrical properties of complex tissue structures. Most biological tissues consist of heterogeneous structures of various cell types that require different stimuli and environments in order to properly function. For example, musculoskeletal tissue is comprised of bone, cartilage, skeletal muscle, tendon, and ligament tissues – all which function upon different cell types, extracellular matrix, stiffnesses, and architectures ^10,11^. Currently, no single bioprinting mechanism can support the multiple networks of soft and hard tissues that are necessary to create viable, functional, and suturable tissue. A combination of bioprinting techniques, however, could potentially assemble multiple soft and rigid materials to engineer more sophisticated and biomimetic tissue constructs with improved properties and functions. Hybrid bioprinting, also called hybprinting, consists of using two or more bioprinting techniques and biomaterials cohesively to print a single construct^12^. By combining multiple techniques and materials, the user can achieve tissue constructs with intrinsic complexities that are more similar to that of native tissue. As of now, current hybrid bioprinting techniques in the literature have mostly worked towards integrating MME and SBM to print rigid polymers within a soft hydrogel network for more physiologically relevant tissue scaffolds^13–18^. In these studies, the hybrid bioprinting of MME and SBM modules was implemented to engineer load-bearing, cellular constructs. The MME module allows for precise extrusion printing of molten thermoplastic material through a high-temperature nozzle to create a layer-by-layer construct. Biomaterials associated with MME, such as poly-caprolactone (PCL) or polylactic acid (PLA), are well-suited for providing sufficient mechanical strength for load-bearing applications and hard tissue engineering, such as musculoskeletal tissue engineering and orthopedics^19–21^. In addition, these biodegradable polymers can be tuned to degrade at a rate that matches native tissue regeneration, making them suitable materials for implant applications in regenerative medicine^22^. On the other hand, SBM allows for feasible extrusion of cells and provides a softer matrix for soft tissue engineering – however it faces issues with low resolution, limited range of materials, and difficulty with support structures during printing^23^. An ideal alternative to SBM is SLA, which enables fabrication of complex architectures with higher resolutions by projection of UV or visible-light high-resolution patterns onto a vat of photocrosslinkable solution. SLA biomaterials, which are typically biocompatible hydrogels, are usually on a softer and more flexible scale than MME, since many SLA materials consist of polymers with elastic properties such as polyethylene glycol-dimethacrylate (PEGDMA), gelatin methacrylate (GelMA), or hyaluronic acid methacrylate (HAMA)^24^. One notable advantage of the photocrosslinkable materials used in SLA is their greater flexibility for material customization through chemical modification of photoreactive groups such as acrylates, methacrylates, or vinyl groups ^25^ – thus, providing a broader range of materials for bioprinting. Prior studies have shown the integration of scaffolds printed via MME with the hydrogel printed with SLA, but typically through multi-step workflows using separate printers ^26,27^. These inter-device transfers require manual handling and re-alignment, increasing risks of contamination, mechanical damage, and dimensional misalignment, while extending fabrication time. To overcome these limitations and leverage the complementary strengths of MME and SLA, we previously introduced the first generation of Hybprinter that executes both modalities within a single automated system^28^. The first-generation prototype, however, was restricted to one material per module and featured limited hardware and software capabilities, and its biological applications were not evaluated.

This study presents an optimized platform for multi-material and multi-cellular hybrid bioprinting to overcome the limitations of using single bioprinting techniques and material. To achieve this, a multi-material hybrid bioprinter, referred to as the Hybprinter, is designed and constructed, incorporating both MME and SLA modules for bioprinting hard and soft tissues. To implement the Hybprinter technology, software components and biomaterials are developed in conjunction with the hardware, enabling automated control of the entire bioprinting process. To expand the functionality of the hybrid bioprinter for multi-tissue bioprinting, a multi-hydrogel bioprinting capability via SLA is integrated into the platform, enabling automated switching between multiple hydrogels and cell types during the printing process. The Hybprinter demonstrated adaptability to multiple soft and hard materials for engineering hybrid constructs that are suturable and implantable. High viability, cell elongation, and proliferation were observed in tissue constructs achieved via SLA and MME/SLA modules of the Hybprinter. Computational and experimental studies were conducted to assess the biocompatibility of the MME technology, specifically investigating the impact of heat generated during MME printing on adjacent cells. Various biological models and therapeutic applications were developed to showcase the broad potential capabilities of the Hybprinter, including limb-on-a-chip, joint-on-a-chip, microfluidics, a vascularized bone model, and a lumbar vertebrae model integrated with an intervertebral disk. Furthermore, stem cell-laden constructs were hybprinted and later differentiated into cartilage and bone tissues to show the Hybprinter’s functionality in multi-tissue engineering.

## 2. Results and Discussion

### 2.1. Hybprinter Development and Construction

The hybrid 3D bioprinter, referred to as the Hybprinter, is constructed and utilized in this study along with its associated fabrication techniques as illustrated in **Figure 1**. The Hybprinter was envisioned for the creation of tissue-imitating constructs that incorporate integrated rigid porous scaffolds and soft hydrogel components. To that end the Hybprinter platform synergistically combines two distinct bioprinting methodologies: Molten Material Extrusion (MME) and Digital Light Processing-based Stereolithography (DLP-SLA), providing an inclusive solution for hybrid tissue bioprinting. The MME module is intended for constructing robust porous scaffolds for structural and mechanical support. The DLP-SLA module, hereafter referred to as SLA, is specifically developed for 3D bioprinting of multi-material soft hydrogels, enabling the spatial patterning and functional integration of distinct hydrogel types that synergistically serve as carriers of different bioactive parts such as cells, drugs, proteins, and nutrients.

**Figure 1.**
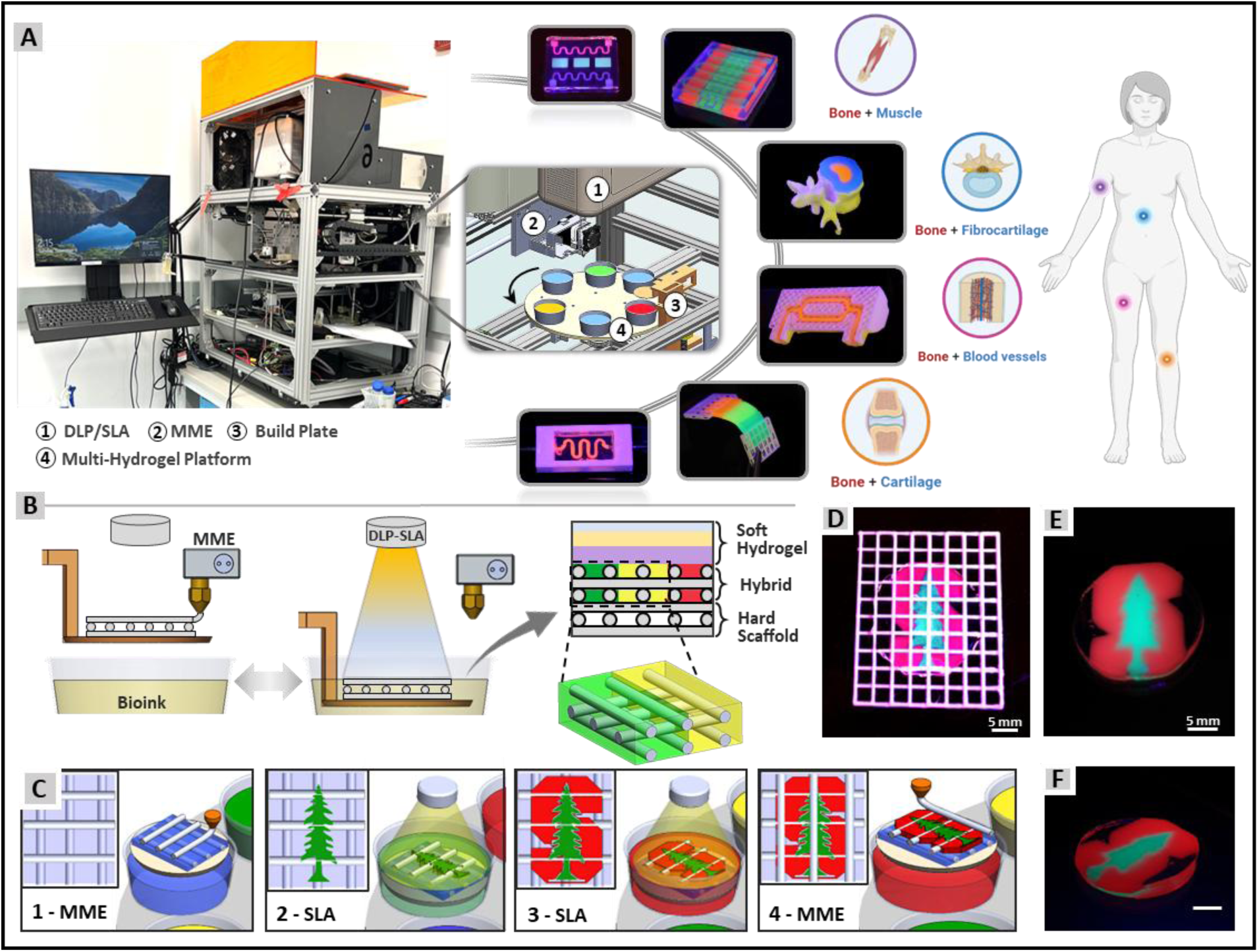
Overview of the 3D hybrid bioprinting approach and applications. (A) Photograph of the Hybprinter with an inset schematic of the printing chamber displaying the MME and multi-hydrogel DLP-SLA modules, alongside sample prints showing potential applications in tissue modeling and therapeutics. (B) Layer-by-layer hybrid fabrication process and schematic representation of a hybrid multi-hydrogel construct. (C) Step-by-step illustration of hybrid bioprinting in progress. (D) Hybrid print showcadsing multi-lydrogel Stanford logo integrated into a hard scaffold. (E, F) Multi-hydrogel sample print of the Stanford logo composed of three colored PEGDMA hydrogels.

The Hybprinter facilitates the engineering of complex constructs that can better mimic multi-tissue interfaces. Biological tissues that vary greatly in stiffness and cellular composition are often interconnected, such as limbs (bones and muscles), vertebra (bones and fibrocartilage), bone marrow (bone and blood vessels), and joints (bone and cartilage). Single module bioprinters are typically limited to printing one type of tissue at a time with a very limited range of stiffness. To address these limitations, multiple demonstration prints were engineered to showcase the potential capabilities and applications of the Hybprinter in fabricating complex tissue engineering constructs, including multi-tissue interfaces, microfluidic models, and multi-cell, multi-hydrogel tissue-mimicking constructs (**Figure 1A**). Each application is discussed in the corresponding segments later in this section.

The hardware components of the hybprinter technology, as depicted in **Figure 1A**, **B** comprise: (1) MME module, which encompasses the extruder, hot end nozzle, temperature control system, as well as the associated electronics, sensors, and controllers; (2) DLP-SLA module, featuring an optical projector; (3) Build plate precise positioning system, with associated electronics, controllers, and sensors; (4) A computer for pre-print model preparation and overseeing the 3D printing process by managing all modules; (5) Temperature control system to maintain the printing chamber at the desired temperature based on the materials used; (6) Multi-hydrogel handling system for incorporating more than one hydrogel in a single 3D bioprinting task. The software aspect of Hybprinter encompasses: 1) Custom-developed pre-print programs for generating printable files for MME and SLA modules, 2) A dedicated program for operating the Hybprinter and executing hybrid printing, and 3) optimized firmware on microcontrollers for hardware control. All programs are developed in MATLAB. The pre-print process involves CAD modeling of the hybrid construct, incorporating both hard and soft components. The CAD model is sliced to generate G-code for the MME module and photomasks for the SLA process, specifying the corresponding materials and vat information. The resulting printing files and instructions are input into the dedicated custom-developed printing program that coordinates the entire printing workflow by controlling the MME microcontroller, SLA projector, positioning system, and multi-vat handling microcontroller.

To fabricate hybrid constructs, a layer-by-layer manufacturing process illustrated in **Figure 1C** is employed, starting from the bottom and progressing to the top on a build plate with precisely controlled motion. For each layer the fabrication begins with deposition of molten material using the MME module and followed by fabrication of the hydrogel component using the DLP-SLA module. The hybrid 3D printing technique is illustrated step-by-step in **Figure 1C**, showcasing the fabrication of the Stanford logo, which comprises multiple hydrogels integrated into a hard scaffold. Additionally, a comparison is made with a version of the Stanford logo printed solely with hydrogels, highlighting the multi-hydrogel printing process. The Hybprinter MME module extrudes molten thermoplastic filaments through a nozzle onto the build plate, following a precisely controlled predefined path to construct a porous mesh structure. The molten thermoplastic material undergoes solidification shortly after extrusion from the nozzle, with solidification being expedited by the implementation of cooling fans attached to the extruder. The MME process is governed by G-code, which is generated during the pre-printing phase through CAD design and subsequent slicing. The hydrogel component associated with each layer is fabricated using the DLP-SLA module. To that end, the build plate is submerged in the pre-polymer solution, where a customized 4K visible-light DLP projector exposes specific regions requiring hydrogel formation to light for a predetermined duration, known as the exposure time, using a projected 2D image referred to as the digital mask or photomask. The photomasks are generated during the pre-printing phase through CAD design and subsequent slicing. Different layer thicknesses can be utilized for MME and SLA modules based on the materials and applications, ranging from 100 to 300 μm. The hybrid construct maintains a default 1:1 integration ratio when each layer of the hybrid construct comprises one layer of MME scaffold and one layer of hydrogel. Nevertheless, the Hybprinter can implement other ratios, such as 1:2 and 1:3, as elaborated in our previous paper ^28^. All materials utilized for the MME and SLA modules, including hard materials and hydrogels, have undergone tuning and testing of their printing parameters; and new materials can also be utilized by tuning the printing parameters and conditions in the software.

The multi-hydrogel printing system shown in **Figure 1A** is integrated into the Hybprinter to enhance its capability to better replicate the architecture of native tissue, incorporating diverse materials and cell types with varying physical, chemical, and biological properties. This multi-hydrogel handling system is an extension of the SLA module, featuring multiple vats, each capable of containing any photocrosslinkable hydrogel, with or without biological agents. The system can accommodate different numbers and sizes of vats, ranging from 25 mL to 95 mL, allowing for a total of 12 to 6 vats, respectively. The actual volume of pre-polymer solution required can be lower than the vat capacity, as the Hybprinter is capable of conducting SLA printing with as low as 5 mL of pre-polymer solution. To prevent cross-contamination, specific vats are designated for washing the build plate when transitioning between different vats. Furthermore, vats can be replaced during the printing process when not actively in use, thereby ensuring operational flexibility. To fully leverage the hybrid capabilities of the system, the SLA multi-hydrogel system can be utilized in conjunction with the MME module to fabricate hybrid multi-hydrogel constructs, as demonstrated in **Figure 1D**, where the Stanford logo, composed of multiple colored PEGDMA hydrogels, is integrated within a hard PLA scaffold. Additionally, the SLA multi-hydrogel system can also function as a standalone platform for printing soft multi-material structures, as shown in **Figure 1E, F**, where the multi-hydrogel Stanford logo is printed. This integration allows for the simultaneous use of hard material and diverse hydrogels to more precisely engineer the intricate hybrid architecture and properties of native tissue.

The bioinks used for SLA printing are composed of a base hydrogel (e.g., GelMA, PEGDMA, HAMA), LAP as the photoinitiator, and tartrazine which acts as the photoabsorber to prevent over-crosslinking and ensure the fidelity of print details^29^. For fabrication of complex constructs having hollow parts, sacrificial bioinks are also utilized to create hollow parts. In this study, two types of sacrifical bioinks are utilized including photocrosslinkable enzymatically-degradable ink (HAMA) and thermoreversible ink (Pluronic F-127).

It should be noted that due to the unique design of the Hybprinter, which integrates two distinct 3D printing modules into a single station, some conventional features of each printing module had to be modified for effective integration. Consequently, the printing parameters and settings typically used for each individual module cannot be directly compared to those employed in this study for the Hybprinter. Specifically, for the DLP-SLA module, standard SLA bioprinters typically employ UV light to accelerate photopolymerization, which can compromise the biocompatibility of bioactive factors and pose a risk for cellular photodamage ^30^. In contrast, the Hybprinter utilizes visible light to maintain cell viability. Additionally, standard SLA bioprinters typically maintain a short distance between the light source and the build plate to leverage higher light intensity for faster photopolymerization and improved resolution. In contrast, the Hybprinter has a significantly greater distance between the DLP-SLA light projector and the build plate due to the presence of the MME module, which also conducts printing on the build plate. Using visible light along with increased projection distance is inherent to the design of the bioprinter and necessitates longer exposure times for photocrosslinking, potentially affecting the achievable resolution compared to conventional SLA technologies. To address these design limitations, a customized high-intensity visible light projector with 3,000 ANSI lumens is integrated into the Hybprinter system, enabling greater penetration depth for more uniform 3D-printed constructs. This light intensity is near the upper end of the typical range for visible light projectors, which operate between 1,500 and 3,500 lumens. A resolution study was performed to determine the minimum line width achievable by the Hybprinter SLA module. The investigation demonstrated that a resolution of 75 μm can be attained (**Figure S3A**), indicating the module’s capability for high-precision sub-millimeter SLA printing. Detailed results are provided in the Supplementary.

### 2.2. Characterization of Hybrid Constructs Properties and Features

One of the primary advantages of hybrid bioprinting is the ability to integrate a hydrogel matrix into a porous hard scaffold, resulting in hybrid constructs. This hybrid functionality is demonstrated in Error! Reference source not found.**A** where a sample square-shaped hybrid construct was 3D printed using PLA as the MME and PEGDMA as the SLA material. The hybrid nature of the fabricated construct was investigated through cross-sectional analysis, which shows that the pores in the hard scaffold are completely filled with hydrogel.

Various materials were incorporated for hybrid bioprinting of MME and SLA modules. For MME, the biocompatible and biodegradable polymers implemented were PLA, PCL, and PCL-β-tricalcium phosphate (PCL/β-TCP)^19^. PLA, PCL and PCL/β-TCP are FDA-approved, clinically available materials commonly used as scaffolds for tissue engineering due to their bioresorbability and mechanical properties^20,21,31–33^. For SLA printing, GelMA, PEGDMA, and hyaluronic acid methacrylate (HAMA) were implemented in the form of photo-crosslinkable solutions. These biocompatible hydrogels enable fabrication of complex high-resolution architectures and constructs by projection of light-based photomasks (**Figure S1-B**). **Figure 2B** demonstrates the wide range of mechanical properties handled by the Hybprinter. For hard materials, a higher range of stiffness is achieved with PCL and PCL/β-TCP at a Young’s modulus of approximately 200 MPa and PLA at around 1300 MPa. In comparison to soft materials, the hydrogels provide the lower range of stiffness required for cells in tissue engineering constructs, with GelMA exhibiting a Young’s modulus of 0.014 MPa and PEGDMA showing a Young’s modulus of 0.05 MPa.

**Figure 2.**
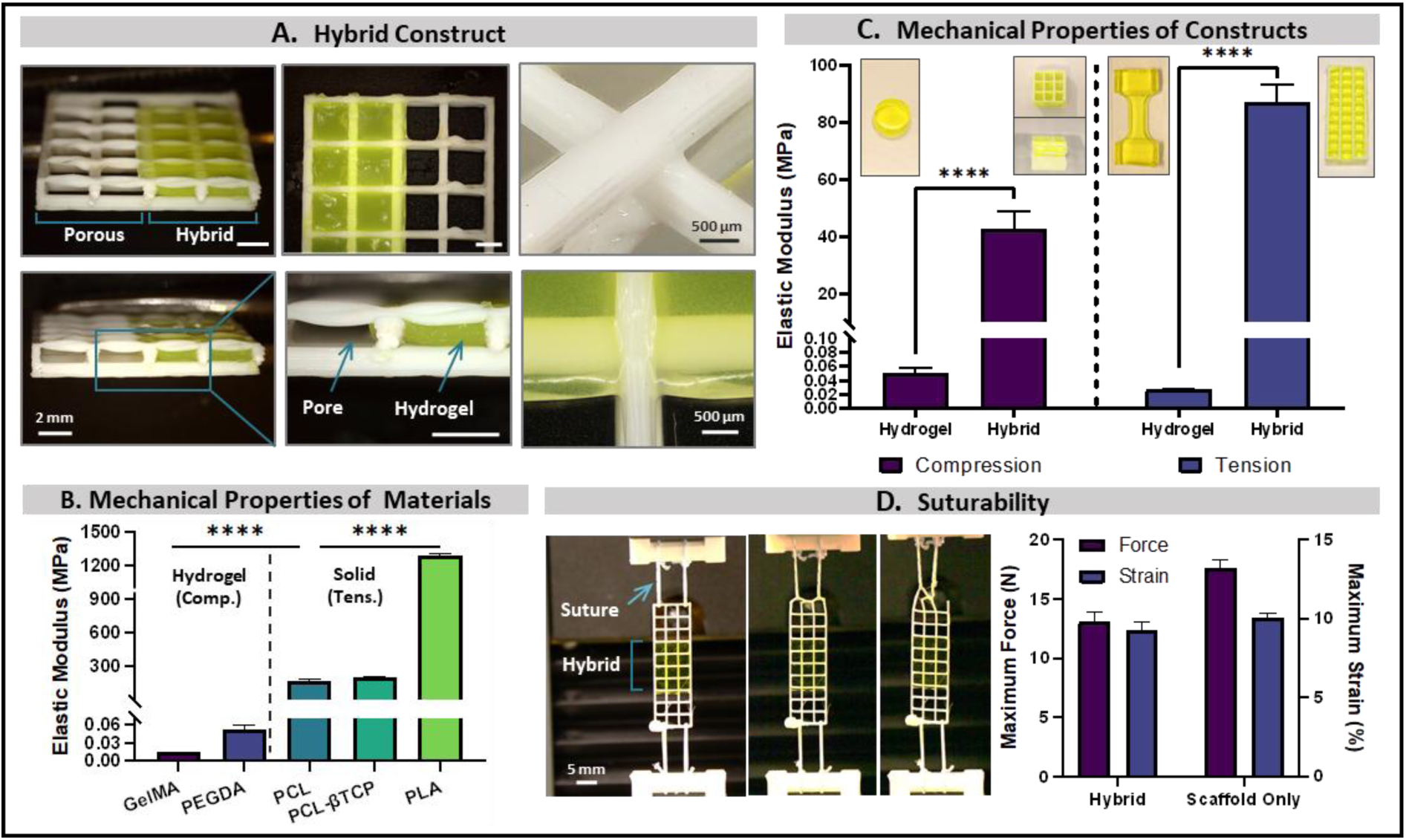
Hybrid construct and material characterization, along with biocompatibility and suturability assessment. (A) Hybrid construct partially integrated with hydrogel, demonstrating complete pore infiltration. Magnified images show junctions between struts on different layers (top) and at the hydrogel interface (bottom). (B) Elastic modulus of materials utilized by the Hybprinter, including hydrogels and solid filaments, under compression and tensile testing, respectively. (C) Mechanical strength of the hybrid constructs compared with similar hydrogel constructs, represented by tensile and compressive elastic moduli. (D) Suture retention test results of hybrid construct and the scaffold without hydrogel.

The mechanical strength of the hybrid construct can be adjusted by two factors: the type of thermoplastic material and porosity. As shown in Error! Reference source not found.**B**, the materials used for Hybprinter can achieve an elastic modulus as low as 0.015 MPa (with GelMA) and as high as 1300 MPa (with PLA). Additionally, the stiffness of the hybrid constructs can be further optimized beyond material selection by controlling pore size during 3D printing via the MME. Porosity can be regulated during the pre-print process, and its critical influence on mechanical stiffness has been systematically investigated in our previous study on biomedical devices and bone tissue engineering^34^. With the versatility to create both soft and/or hard constructs, the Hybprinter technology allows for 3D-printed constructs that can achieve a wide range of stiffnesses that otherwise wouldn’t be feasible with printers that only perform single bioprinting techniques.

Mechanical testing of hybrid bioprinted constructs demonstrates an important solution offered by Hybprinter to engineer enhanced mechanical strength into 3D constructs, as illustrated in Error! Reference source not found.**C**. Hydrogels alone lack sufficient strength to provide mechanical support by showing tensile and compressive modulus less than 0.1 MPa, making them unsuitable for load-bearing medical implants, particularly in musculoskeletal tissue engineering and also for supporting sutures, however, as shown in Error! Reference source not found.**C** when PLA is hybrid printed within a hydrogel, the resulting construct demonstrates a 1000-fold increase in compressive strength and a 4000-fold increase in tensile strength compared to hydrogel alone. In comparing the implementation of a non-load-bearing hydrogel to a hybrid hydrogel, it is evident that while both share similar gel and biological properties, the hybrid construct offers significantly greater mechanical strength. This capability is a unique advantage of the Hybprinter technology, as most other bioprinters cannot achieve such a combination of flexibility and strength. The ability to create cellular constructs that not only mimic the biological environment but also provide the necessary mechanical support is crucial for advancing applications in tissue engineering and regenerative medicine.

Bioprinting of a suture-compatible tissue-engineered construct may appear straightforward; however, most current bioprinters that support cellular encapsulation produce constructs that cannot be sutured, presenting a significant challenge for tissue implant development^35^. The Hybprinter overcomes this limitation by seamlessly integrating hydrogel and hard filament to create hybrid constructs that facilitate both cell encapsulation and suturing. The suturability of these hybrid constructs was assessed through a suture retention test, with results shown in Error! Reference source not found.**D**. The experiment demonstrated that the hybrid construct, with 85% porosity, can withstand up to 13 N of tensile force and 9% maximum tensile strain.

Although the wet condition of the hybrid scaffold slightly reduces mechanical strength compared to a same dry non-hybrid scaffold, the force levels remain sufficiently high to allow for effective suturing. The suture retention strength discussed here refers to the maximum force that can be applied to the suture thread before implant failure or suture migration occurs. In this study, this metric is intended for comparison among hydrogel-based bioactive grafts and patches, which serve as practical reference points for implant applications, and should not be compared to native tissues or nonresorbable patches, as the focus here is on suturable hydrogel-based tissue-engineered constructs. Currently available solutions, such as decellularized collagen patches and sponges capable of carrying cells and sutur retention, are not hydrogel-based^36^. Utilizing the Hybprinter, the suture retention strength of hybrid scaffolds can be optimized for specific applications by selecting from various scaffold materials and adjusting scaffold porosity.

### 2.3. Hybprinter Biocompatibility

To validate the hybprinter technology as a bioprinter that supports both viability and proliferation for tissue engineering applications, the SLA module was first validated by 3D bioprinting GelMA constructs encapsulated with bone marrow-derived, human mesenchymal stem cells (hMSCs). Firstly, the light exposure time was tuned in response to hMSC viability and GelMA hydrogel stiffness by also respecting the printing fidelity. Results show that high cell viability was achieved with 90 seconds of exposure time (per layer) on days 1 and 3 after printing with over 90% viability achieved on day 3 (**Figure S3**)^37^. As a result, the 90-second time interval for SLA bioprinting was determined as the near-optimal exposure time in terms of cell viability and hydrogels stiffness that supports printing fidelity and was implemented for all further experiments detailed here unless stated otherwise.

The behavior of the cell-laden constructs fabricated with Hybprinter is investigated over a two-week period by evaluating the viability and proliferation of SLA-bioprinted hMSCs in GelMA. Viability starts at around 80% one day after printing, then reaches around 90% after two weeks, suggesting viable cells are proliferating during that time-frame (Error! Reference source not found.**A, B**). On Day 7, hMSCs start to elongate before forming cellular networks by Day 14. This is further supported by quantification of hMSC proliferation, as cells appear to increase proliferation rates following each timepoint (Error! Reference source not found.**C**). This evidence suggests that the SLA module of the Hybprinter supports stem cell viability, proliferation, and elongation that are crucial for creating functional tissue-like constructs – making it a suitable platform for tissue engineering and stem cell therapy applications. Beyond cell viability and proliferation, differentiation of stem cells in bioprinted constructs was investigated to explore the Hybprinter’s capability in tissue engineering, as discussed in the corresponding parts later in this section. Moreover, during vat photopolymerization of cell-encapsulated hydrogels, variations in cell density within layers due to gravity and sedimentation can raise concerns regarding nonuniform cell distribution. In multiple samples of cell-laden GelMA constructs with 300 μm thickness per layer, Error! Reference source not found.**D** demonstrates uniform cell distribution across layers, as evidenced by normalized standard deviations (i.e., coefficients of variation) of cell densities among layers in different samples, all of which are considerably below 10%. As a result, all further cellular bioprints discussed here are performed using these optimized parameters.

**Figure 3.**
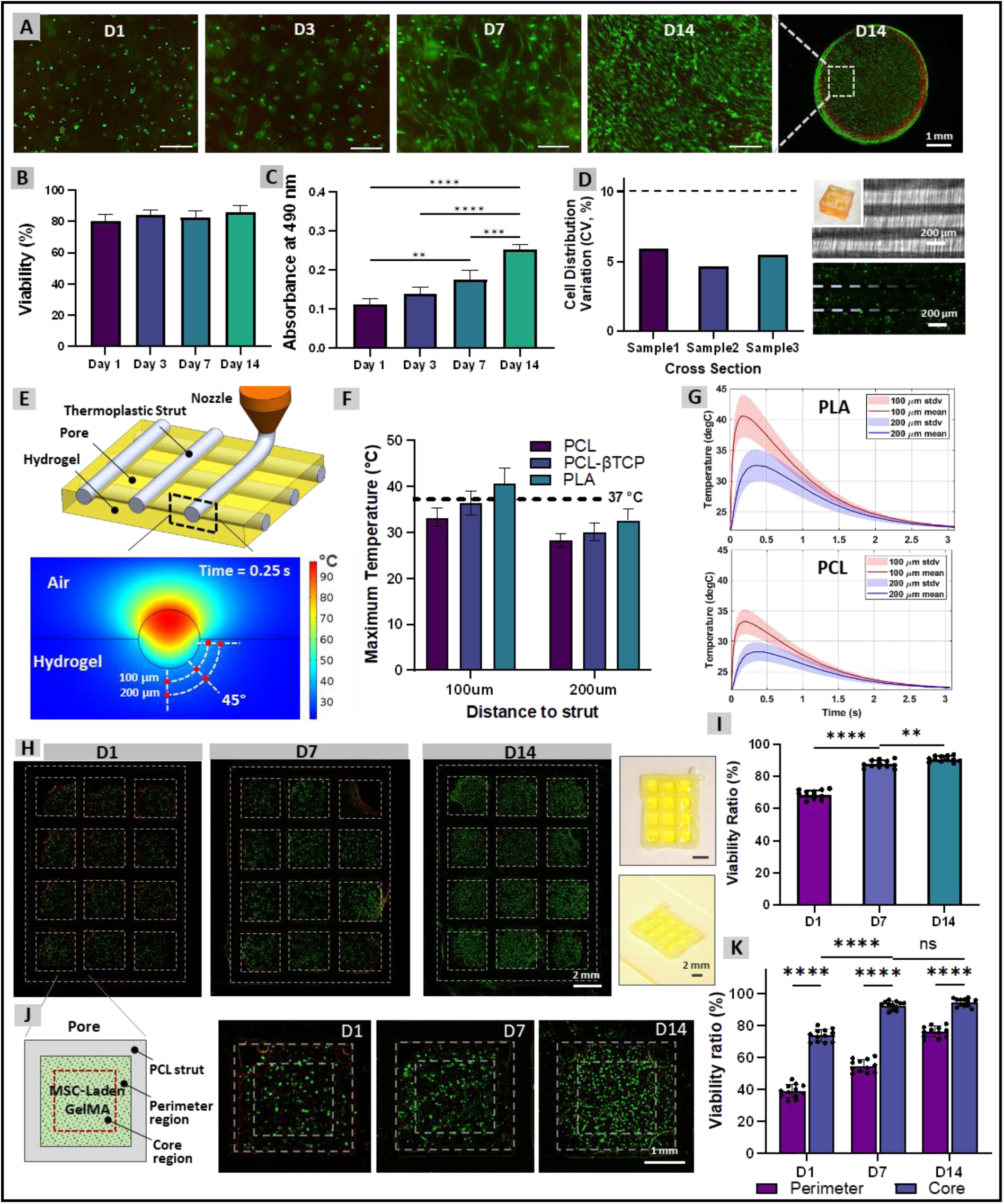
Biocompatibility of the hybrid bioprinting approach and viabilty of the farbricated hybrid constructs. (A) Live-dead fluorescent imaging of MSC-laden GelMA hydrogels over 14 days. (B) Viability ratio of the cellular prints. (C) Proliferation quantification of the cell-laden prints. (D) Cell distribution uniformity across SLA print layers, assessed by fluorescent imaging and quantified by the coefficient of variation. (E) Schematic of MME printing as part of the hybrid bioprinting process, illustrating the computational thermal modeling domain and a simulation snapshot showing the temperature distribution, circular MME-printed strut, hydrogel, and temperature measurement probes. (F) Maximum hydrogel temperature predicted by the computational model at designated measurement points for PCL, PCL-TCP, and PLA filaments, printed at 130°C, 160°C, and 200°C, respectively. (G) Temporal variation of hydrogel temperature obtained from computational simulations. (H) Live-dead fluorescent imaging of MSC-laden hybrid constructs over 14 days, along with a photograph of the printed construct. (I) Viability ratio of the hybrid constructs over the 14-day period. (J) Schematic of a single hydrogel pore, highlighting the perimeter region adjacent to the scaffold strut and the core region, with corresponding live-dead fluorescent images. (K) Viability assessment of cells in the perimeter versus the core region, over 14 days.

Following the demonstration of biocompatibility and optimization of hydrogel SLA printing, it is crucial to investigate and validate the biocompatibility of the Hybprinter in fabrication of bioactive hybrid constructs that involve MME, SLA, and cell-laden bioinks. This investigation is conducted through both computational and experimental approaches. A primary concern regarding the incorporation of MME in hybrid bioprinting is the effect of heat generated by the extruded thermoplastic materials on cell viability. To address this, a computational thermal simulation was first performed to assess the impact of MME within a hydrogel platform in real-time (Error! Reference source not found.**E-G**). The results from computational modelling and simulation suggest that the immediate post-print temperature rise in the hydrogel, caused by the hot strut, does not exceed 37°C and is confined within a minor zone around the strut less than 200 μm. It is critical to note that bioprinting parameters maintaining thermal conditions at or below 37°C are typically considered optimal to preserve cell viability. Even closer to the strut at a proximity of 100 μm, only the PLA material model induces a brief temperature rise within the hydrogel above 37°C, lasting less than one second. Moreover, the hybrid construct’s substantial porosity, typically greater than 85%, results in wide strut-to-strut distances, yielding pore sizes larger than 3 mm. Consequently, the affected radius around the strut, within 200 μm, is negligible and does not adversely affect the viability of most of the cells distributed within the pore spaces. It should be acknowledged that hot thermoplastic materials may impact cells in direct contact with the hot strut during printing, although such cells represent a minor proportion relative to the large population of cells encapsulated inside the hydrogel within the pores. Therefore, within the hybrid bioprinting context, hot thermoplastic materials do not undermine the viability of cells integrated within the hybrid construct.

An experimental approach was undertaken in conjunction with the computational approach to validate the biocompatibility of the Hybprinter and confirm the computational results. To this end, cell-laden hybrid constructs were 3D printed, incorporating hMSCs within a GelMA bioink via SLA, alongside a PCL porous scaffold fabricated using MME. The viability of the hybrid constructs was assessed over a 14-day period. As shown in Error! Reference source not found.**H-I**, around 70% cell viability was achieved Day 1 after printing, as cells near the PCL struts particularly appear to be impacted. By Day 7, however, cells appeared to have recovered and maintained high viability (∼90%) up until Day 14. To further investigate the effect of MME on cell viability, the viability of the cells printed closest to the PCL strut (1/6th of the pore width, labeled as the perimeter) versus the cells printed the farthest from the PCL strut (labeled as the core). Error! Reference source not found.**J-K** shows a viability difference of about 47% observed in Day 1 between the perimeter and core of the pores, however this difference reduces to less than 20% by Day 14. In addition, cell spreading and elongation is observed by Day 14, which serves as a positive indicator of biological function in hMSCs within the hybrid constructs^38^. This evidence further supports our hypothesis that cells that are printed closest to the PCL strut are initially impacted by MME but manage to recover and maintain high viability long-term. These results demonstrate a hybrid bioprinter technology that is biocompatible and promotes cell elongation of stem cells.

### 2.4. Multi-Material Hybrid Bioprinting and Applications

This section outlines the potential applications of the Hybprinter, following the preceding discussion on technology, biomaterials, and biocompatibility. In general, the majority of the bioprinting efforts are focused on one of two primary applications: (1) engineering representative in vitro models and (2) recreating biological constructs and implants for therapeutic purposes. However, single-module bioprinting systems are often restricted in terms of biomaterials, the number of cell types, and the range of achievable stiffnesses. As a result, their ability to recreate the complexity of native tissues and their functions is significantly reduced. By combining multiple bioprinting techniques and materials into one system, the Hybprinter expands the range of biological tissues that can be fabricated compared to single-module bioprinting. This section explores some modeling and therapeutic applications enabled by the Hybprinter, acknowledging that the potential applications extend beyond those discussed, with selected examples addressing key ongoing challenges that can uniquely solvable by the Hybprinter.

#### 2.4.1 Microfluidics and Organ-on-a-chip

Microfluidic devices are increasingly utilized for engineering organ-on-a-chip platforms to model human physiology and pathology due to their customizable, user-friendly, and cost-effective nature^39,40^. Developing these platforms typically involves fabricating a microfluidic device, sterilizing it, and seeding cells or other relevant factors to mimic an organ. Conventional fabrication processes, such as soft lithography, consist of multiple steps for microfluidic device assembly which can span over several days before reaching a finalized organ-on-a-chip model^41^. In comparison, 3D printing has allowed for rapid prototyping of microfluidic devices with significantly lower labor costs that can then be sterilized and seeded with growth factors and cells for microfluidic purposes – but like soft lithography, traditional 3D printing methods are mostly limited to molding devices that later must be sterilized and prepped for cellular integration^42–44^. Due to the Hybprinter’s top-down SLA design, microfluidic devices can be fabricated with sterile-filtered inks along with cell-laden hydrogels in one streamlined process, thus automating the organ-on-a-chip fabrication, sterilization, and cell-seeding process. Error! Reference source not found.**A** demonstrates the multiple microfluidic channel orientations achievable with the Hybprinter, both throughout the x- or y-plane or throughout the z-plane (cell-laden hydrogels and microfluidic channels represented by color). Moreover, for load bearing tissue models, the integration of multipe hydrogels into a hybrid construct is exemplified in Error! Reference source not found.**B**, demonstrating the side-by-side integration of biomaterials in a hybrid configuration. This hybrid bioprinting approach enables automation of microfluidic device fabrication and organ-on-a-chip assembly within a single system in a relatively short duration, positioning the Hybprinter as a promising platform for rapid prototyping.

**Figure 4.**
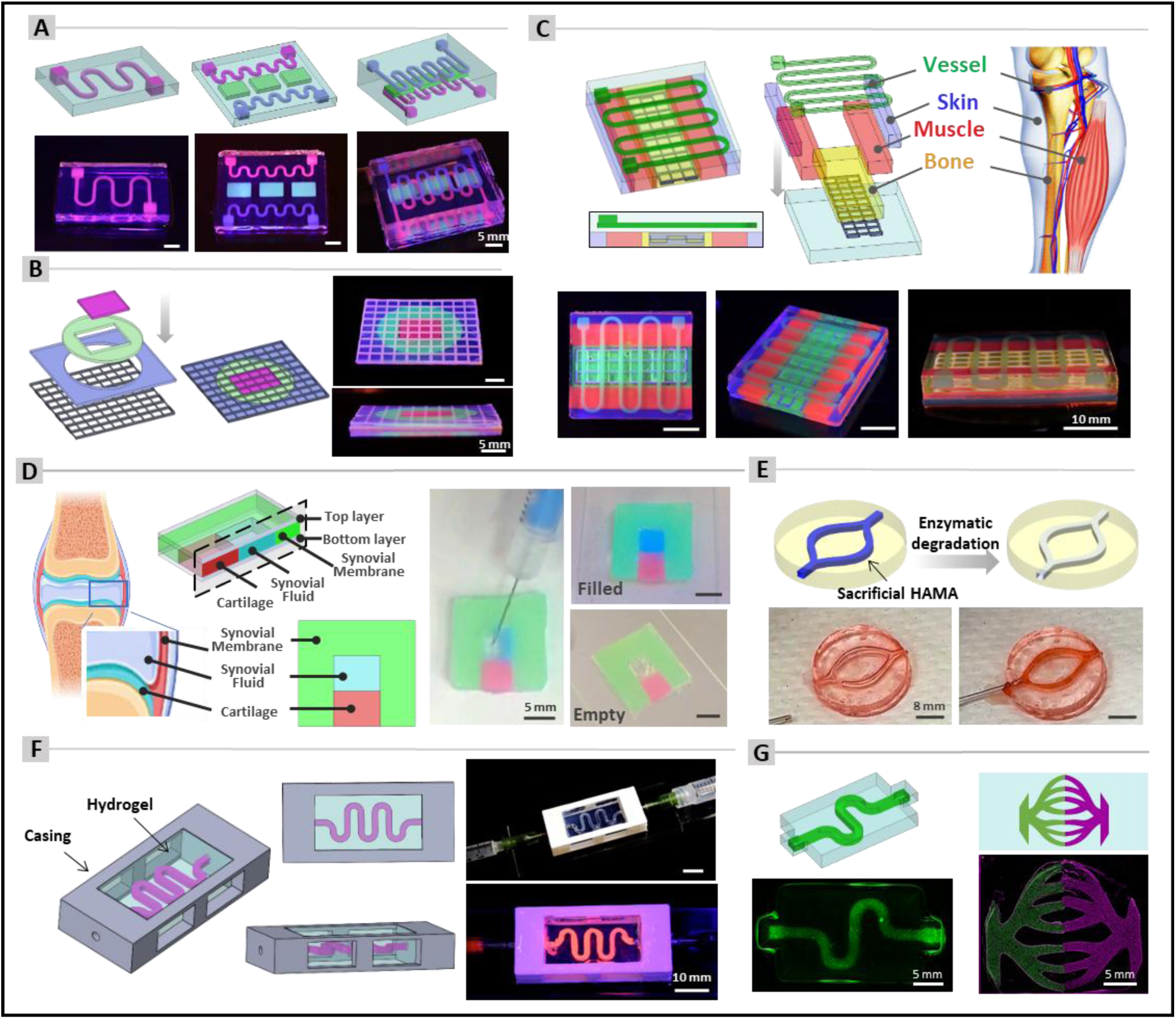
Hybprinter potential applications for tissue modeling. (A) Representative multi-hydrogel prints for microfluidic and organ-on-a-chip applications. (B) Hybrid multi-material model for load bearing interfacial tissue modelling. (C) Limb-on-a-chip featuring a central hybrid section representing bone and two side regions representing muscle and skin, with an integrated channel for vascularization. (D) Joint model illustrating cartilage and synovial membrane, including a hollow internal cavity for injecting and retaining synovial fluid while preventing mixing with culture media. (E) Perfusable construct for vascularization, produced by enzymatic degradation of sacrificial HAMA. (F) Robust perfusable construct integrated with a casing, fabricated simultaneously in a single printing job. (G) Cellular multi-material prints utilizing HUVECs tagged with different fluorescent markers for vascularization modelling.

#### 2.4.2 Limb-on-a-chip

Modeling vascularized musculoskeletal tissues within an in vitro platform remains challenging due to their broad stiffness spectrum. Single-module bioprinters cannot achieve both soft musculoskeletal tissues (such as muscles, tendons, and cartilage) and hard musculoskeletal tissues (such as bone) within a single construct – thus, leaving a need for more advanced bioprinting technologies. By combining the fabrication of microfluidic and multi-tissue platforms, the Hybprinter system is capable of engineering multi-tissue-on-a-chip technologies. Error! Reference source not found.**C** demonstrates a skin-muscle-bone-vascular model on a chip (porous scaffold + yellow = bone, red = muscle, purple = skin, green = vasculature), where the bone portion is hybrid and the other tissues are hydrogel-based. In addition, the Hybprinter can further tune hydrogel stiffness and filament porosity to mimic the stiffness range between these tissues while also providing support to carry each tissue’s corresponding cells and/or bioactive cues. By recreating these complex, multi-tissue interfaces in a single construct, the Hybprinter lays the foundation for next-generation in vitro models that can more accurately recapitulate musculoskeletal physiology.

#### 2.4.3 Joint model

Bioprinting a joint-mimicking model presents challenges due to the intricate architecture of multiple tissue types, including bone and cartilage, alongside the synovial fluid encapsulated in the synovial membrane lining of synovial joint cavities. The synovial fluid, although present in minimal volumes, is essential for joint biological function and development. The Hybprinter was utilized to fabricate a joint model that integrates synovial fluid within the synovial membrane, in conjunction with cartilage tissue. Fabrication of a hollow internal cavity that can hold fluid within a multi-hydrogel structure presented a bioprinting challenge, which was addressed in this model using sacrificial materials. This approach aims to replicate the native joint environment more accurately, facilitating improved biological interactions and mechanical stimulation. The development of this model is guided by two primary objectives: the ability to facilitate diffusion toward the hollow cavity and the capacity for mechanical stimulation. The first objective is achieved through the fabrication of a diffusible multi-hydrogel construct that can be cell-laden, incorporating a hollow structure encased within the model containing synovial fluid, as illustrated in Error! Reference source not found.**D**. This model ensures the separation of synovial fluid from the culture media while facilitating the diffusion of the culture media throughout the entire construct. This diffusion process is critical for delivering essential nutrients, oxygen, and necessary elements for cellular viability. The secondary objective of enabling mechanical stimulation is achieved through the inherent elastic properties of the hydrogel, which comprise the entire model. These properties facilitate deformation under mechanical forces, which are essential for promoting cellular responses and tissue development in musculoskeletal tissue engineering and mechanobiology studies. In contrast, conventional tissue-on-a-chip models constructed from polydimethylsiloxane (PDMS) lack the capacity for mechanical stimulation, limiting their applicability in dynamic loading environments. This model was fabricated using thermoreversible sacrificial Pluronic which is extruded manually via syringe to preserve the hollow region for synovial fluid during printing. The sacrificial part was subsequently liquefied and removed at a lower temperature, enabling the injection of a fluid that simulates synovial fluid, as shown in Error! Reference source not found.**D**. This approach demonstrates the potential application of the Hybprinter in developing diffusible, flexible models that incorporate enclosed hollow structures.

#### 2.4.4 Vascularized models

One of the main obstacles in tissue engineering and bioprinting is recreating a tissue’s vascular system. Engineering perfusable and mechanically stable channels within a 3D-printed construct can be achieved through embedded bioprinting (via sacrificial ink) or inverted SLA methods. Embedded bioprinting can achieve perfusable channels within a hydrogel construct using sacrificial ink but the resolution is quite low, whereas inverted SLA can achieve high-resolution prints of perfusable channels without sacrificial ink. In both cases, however, the handling and installation of a perfusion flow system within these printed constructs remains challenging due to the soft nature of the hydrogel. The Hybprinter’s multi-hydrogel printing capability allows for the use of sacrificial bioinks, enabling the precise fabrication of perfusable channels within hydrogel constructs. In the first demonstration shown in Error! Reference source not found.**E**, HAMA is employed as a suitable sacrificial ink that can be photocrosslinked and subsequently degraded enzymatically with hyaluronidase. Additionally, HAMA is biocompatible, which can allow for bioprinting cell-coated, perfusable channels within constructs^45^. In this model, a perfusable channel was created using 1% HAMA ink, which was subsequently degraded enzymatically by hyaluronidase. Particularly for cellular bioprinting, the use of hyaluronidase is ideal due to its biocompatibility and lack of interference with GelMA. The second demonstration, illustrated in Error! Reference source not found.**F**, showcases the incorporation of hard materials, highlighting the Hybprinter’s innovative method for fabricating integrated perfusable hydrogel constructs within a hard casing in a single print job. While the creation and assembly of a hard casing surrounding a soft perfusable construct is not inherently complex, the concurrent production of both components during a unified bioprinting process results in a pre-assembled construct that effectively addresses longstanding challenges such as minimizing potential damage during assembly and handling, sealing leakage at inlets and outlets, and reducing contamination risk. In this model the channel is created using sacrificial Pluronic. The side openings of the hard casing facilitate the SLA fabrication process and enhance capillary formation toward the outer regions of the construct, thereby promoting vascularization.

The implementation of endothelial cells in bioprinting for vascularization is investigated by Hybprinter. The multi-hydrogel bioprinting allows for 3D bioprinting of multiple cell-types and materials within a single device, as shown in Error! Reference source not found.**G** where HUVECs tagged with different fluorescent markers were printed within GelMA and PEGDMA gels. Error! Reference source not found.**G** (left) showcases a printed cell-laden microchannel structure within an acellular PEGDMA body. This validated technique^45^ is particularly beneficial for vascularization of bioprinted constructs by establishing HUVEC-laden channels that provide suitable conditions for cell seeding and attachment to the channel walls, followed by non-cytotoxic enzymatic degradation of the hydrogel channel. Error! Reference source not found.**G** (right) presents a more anatomically relevant representation of this vascularization strategy, illustrating two distinct cell lines that model capillaries connecting arteries and veins. Ultimately, the Hybprinter serves as a valuable tool for developing advanced vascularized models to study the perfusion flow of biological tissues.

In the realm of tissue modeling discussed thus far, the Hybprinter’s integrated approach – combining MME, SLA, and multi-material bioprinting – facilitates the rapid prototyping of multi-cellular and multi-hydrogel microfluidic devices, organ-on-a-chip models, and vascularized constructs, thereby enhancing the fidelity of native tissue mimicry. Transitioning to therapeutic applications, the Hybprinter exhibits notable applicability in the field of therapeutics as illustrated by the following examples including multi-tissue interfaces, vascularized bone grafts, and hybrid vertebrae-disc replacement grafts.

#### 2.4.5 Multi-tissue interfaces

For therapeutic applications, Hybprinter allows for automated fabrication of multi-material hybrid constructs that can address the requirements for interfacial tissues with load-bearing capacity, such as musculoskeletal interfacial tissues including bone-tendon, bone-ligament, and bone-cartilage junction sites^46,47^. Single-module bioprinters are usually limited to a small range of stiffness, biomaterials, and one to two extruders or vats for multi-cellular printing – hence, making it difficult to study biological tissues which consist of a wide range of stiffnesses, microarchitectures, and cellular compositions. Ligaments and tendons including interfacial fibrocartilage tissue, for example, are soft tissues that range in tissue composition and stiffness which is essential for providing structural support and connection between bones or bone and muscle. Multi-hydrogel hybrid bioprinting enables the 3D modeling of ligaments (Error! Reference source not found.**A, left**) and fibrocartilaginous tendons (Error! Reference source not found.**A, right**) by incorporating varying tissue compositions and stiffnesses along the x, y, and z axes, i.e. 3D (Error! Reference source not found.**A**). The resulting hybrid constructs present a potential platform for more physiologically relevant implants and tools for studying mechanical stimulation and mechanobiology of multi-tissue interfaces.

**Figure 5.**
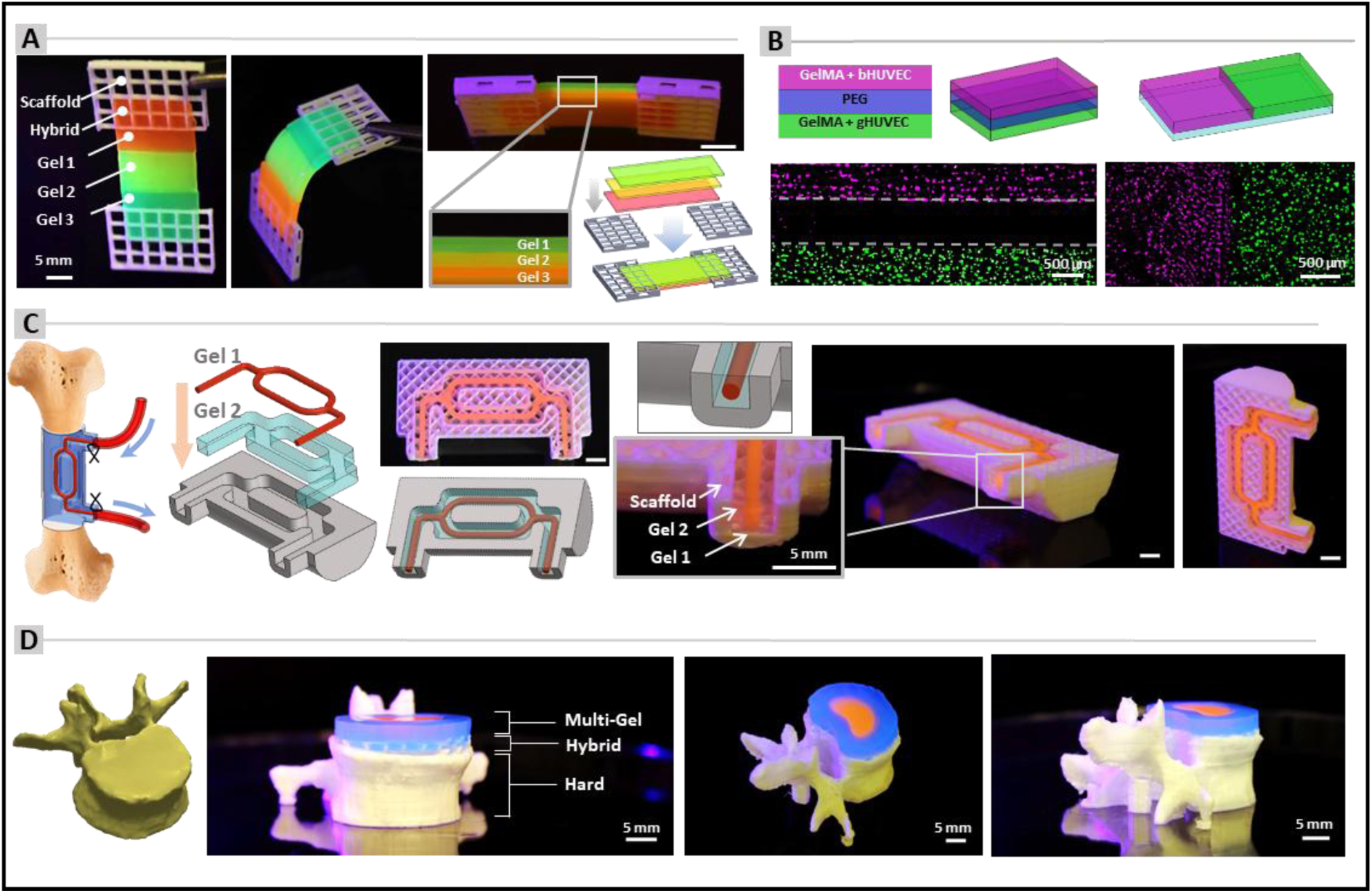
Hybprinter potantial applications in thraputics. (A) Representative hybrid multi-hydrogel prints for interfacial multi-tissue engineering. (B) Multi-cellular, multi-material prints demonstrating multi-layer interfacial tissue engineering. (C) Vascularized bone graft with load-bearing capability and an integrated channel suturable at both the inlet and outlet. (D) Hybrid vertebra–disc replacement and model.

The simultaneous incorporation of multiple cell types and hydrogel materials within a single printed construct is demonstrated through the fabrication of biologically relevant cellular constructs utilizing two different hydrogels (GelMA and PEGDMA) and two different cell lines (bHUVEC and gHUVEC). This results in three distinct bioinks, with GelMA serving as the base hydrogel for both cell types and PEGDMA functioning as an acellular bioink. Error! Reference source not found.**B** (left) illustrates a three-layer construct comprising distinct cellular top and bottom layers with an acellular middle layer, applicable for the repair and correction of multi-tissue defects. Error! Reference source not found.**B** (right) illustrates two side-by-side multi-cell constructs positioned on an acellular substrate layer.

#### 2.4.6 Vascularized bone graft

In clinical practice, bone grafts are either derived from the patient’s own bones (autografts), cadaveric bone (allografts), or fabricated from synthetic materials (synthetic grafts). Patients, however, have limited bone tissue that can be harvested and grafted to other areas, allografts carry risks of immune rejection and disease transmission, and synthetic hard materials cannot adequately replace large bone defects. Previous bioprinting studies have shown the use of thermoplastic materials as bone implants for treatment of large-bone defects and while these rigid materials can offer the structural and load-bearing features necessary to support bone properties, they lack the proper cell-carrier system to offer biological support for enhanced bone regeneration. Although classical MME printing, when used alone, allows for post-fabrication cell loading to prevent heat-induced cell damage, it still lacks the spatial control of cells within the resulting constructs. The Hybprinter’s capability to combine both soft and hard tissue constructs can be advantageous in achieving a vascularized bone graft. In Error! Reference source not found.**C**, a schematic shows the anatomical inspiration of a vascularized long bone, in which a principal vessel travsers the graft to ensure blood supply. Using the Hybprinter, the graft design integrates a porous, mechanically robust, hard scaffold for structural and mechanical support and a dual hydrogel-based channel system – one serving as a major perfusable vascular conduit (Gel 1) and the other as a cell carrier system (Gel 2), to promote vascularization and graft integration. In the printed bone graft construct, the hard scaffold is designed with interconnected pores that facilitate bone ingrowth, while the integrated perfusable main channel facilitates nutrient and oxygen transport, as well as capillary formation. Additionally, the integration of coaxial hard material at the inlet and outlet with a soft hydrogel channel holds promise in effectively mitigating suturability concerns associated with major blood vessels. Overall, this model illustrates that the Hybprinter could facilitate the production of multi-material, vascularized bone graft with the structural strength, suturability, and biological properties necessary for the treatment of large bone defects. Furthermore, while this model incorporated essential features of a hybrid bone graft, the Hybprinter enabled the fabrication of more advanced constructs in which the pores of the hard scaffold, representing bone, were filled with cell-laden hydrogel, potentially accelerating the bone healing process.

#### 2.4.7 Hybrid vertebrae-disc replacement graft and model

Full or partial vertebral body replacement is an optimal corrective intervention for patients with severe spinal trauma, tumors, or infections. Spinal reconstruction with vertebral body replacements usually consists of metal fixation systems that provides the structure and load-bearing support necessary to replace the vertebral body – however, it severely impacts the motor function of patients due to soft tissue damage of adjacent discs^48,49^. Intervertebral disc replacements aim to treat damaged spinal disks and can help improve motor function of the spine, but engineering a replacement with the hard tissue properties of a vertebrae in conjunction with the soft tissue properties of an intervertebral disc remains challenging due to the vast difference in mechanical needs. The Hybprinter has the potential to combine both hard and soft tissue printing to engineer a vertebral body replacement with an integrated artificial intervertebral disc, as modeled in Error! Reference source not found.**D**. Utilizing Hybprinter, a patient specific L2 lumbar vertebral body was 3D printed via MME using a hard scaffold where the top portion is seamlessly integrated with an artificial disc 3D printed via MME and SLA in a hybrid scaffold. As illustrated in Error! Reference source not found.**D**, the printed construct comprises a hard component (representing the vertebrae), a hybrid section (representing the interface between the vertebrae and disc), and a multi-hydrogel region (representing the intervertebral disc). With Hybprinter, the hydrogel portion can also be printed with embedded cells, enabling both modeling and therapeutic applications for spinal injuries. As a potential implant, the hybrid vertebrae-disc replacement may reduce adjacent disc degeneration and preserve motor function compared to conventional vertebral body replacements.

### 2.5. Chondrogenic and Osteogenic Differentiation in Hybprinter Constructs

The objective of bioprinting is to fabricate constructs that closely mimic multi-tissues and their interfaces by implementing a wide range of biomaterials, stiffness variations, and bioprinting techniques. To achieve this with Hybprinter, in addition to assessing biocompatibility, it is thus crucial to investigate whether the Hybprinter can achieve cellular constructs that support stem cell differentiation for multiple tissue types. Here, hMSC-laden constructs were bioprinted and underwent chondrogenic and osteogenic differentiation. For cartilage tissue engineering, hMSC-laden gel (GelMA) constructs were bioprinted via SLA to mimic softer tissue and then underwent chondrogenic differentiation for 28 days (Error! Reference source not found.**A**). To bioprint multiple samples during a single print run, a photomask of a network of 4-mm disks was implemented (**Figure S1B**). The branched network between the disks supports efficient bioprinting of multiple samples by eliminating the risk of sample loss during printing. Immediately after printing, disks were separated into individual wells within a 24-well plate; henceforth, microscopy images of gel constructs may show one to three branched structures near the perimeter. To confirm the presence of chondrocytes, histology and immunostaining methods were performed after 14 and 28 days of differentiation to observe potential key components of extracellular matrix (ECM) in cartilage. As such, the presence of glycosaminoglycans (GAGs) was confirmed via Alcian Blue staining after 14 and 28 days of differentiation, with a stronger signal observed in the latter timepoint (Error! Reference source not found.**B**). Due to the hydrogel’s absorption of the dye, a light-blue tint can be observed in the background and thus an acellular control group that was stained side-by-side with the differentiated group is displayed for fair comparison. Furthermore, it is hypothesized that due to water absorption and gradual hydrogel degradation during the incubation period, the Day 28 control group appears larger than Day 14. However, the differentiated constructs in Day 28 appear smaller than the controls, which we hypothesize is a result of contraction by cells and cellular deposition of cartilage matrix^50–53^. In addition, constructs were stained for collagen type II (COL-II) which is the most abundant protein found in native articular cartilage tissue (Error! Reference source not found.**C**). A negligible level of COL-II was observed on Day 14, but by Day 28, constructs demonstrated a high abundance of COL-II expression. The confirmation of two essential components of cartilage ECM, GAGs and COL-II, demonstrates the Hybprinter’s capacity for cartilage tissue engineering.

For bone tissue engineering, hMSC-laden gel (GelMA) and hybrid (GelMA/PCL/β-TCP) constructs were bioprinted via SLA and MME + SLA, respectively. In addition to the enhanced osteogenic potential provided by β-TCP ^54–56^, the hybrid construct can allow for a stiffer environment that is more representative to bone than gel alone, which can allow MSCs to differentiate into osteogenic lineage more robustly^57–59^. Thus, to investigate the potential benefits of bone tissue engineering using Hybprinter methods, both hybrid and gel constructs underwent osteogenic differentiation for 28 days.

To characterize the osteogenic potential of the gel and hybrid constructs, alizarin red staining (ARS), immunostaining, and micro-computed tomography (micro-CT) were performed after 21 and 28 days of osteogenic differentiation (**Figure 6**). Due to the intense color of the ARS in the constructs, imaging settings were adjusted accordingly per sample to improve visibility of the staining. The hybrid constructs demonstrated darker and larger calcium deposits in comparison to gel constructs for both timepoints (**Figure 6D-E**). On Day 21, in the Gel group, ARS revealed scattered mineralized nodules (red arrows), indicating the onset of calcium deposition. In contrast, the Hybrid group exhibited more uniform and denser ARS, suggesting earlier and more extensive mineralization compared to the Gel group. Also, the gel portion of the hybrid constructs underwent more contraction than the gel constructs, most likely due to the hybrid’s stiffer environment^59^. By Day 28, all constructs appear to have higher water intake due to the longer incubation period and thus, cell-induced contraction is not as apparent as in Day 21. Calcium deposits are more apparent on Day 28 for both gel and hybrid constructs. In the Gel group, mineral deposition increased relative to Day 21, with larger clusters of mineralized nodules (white arrows). The Hybrid group, however, showed consistently strong and widespread ARS, reflecting sustained and more mature mineralization.

**Figure 6.**
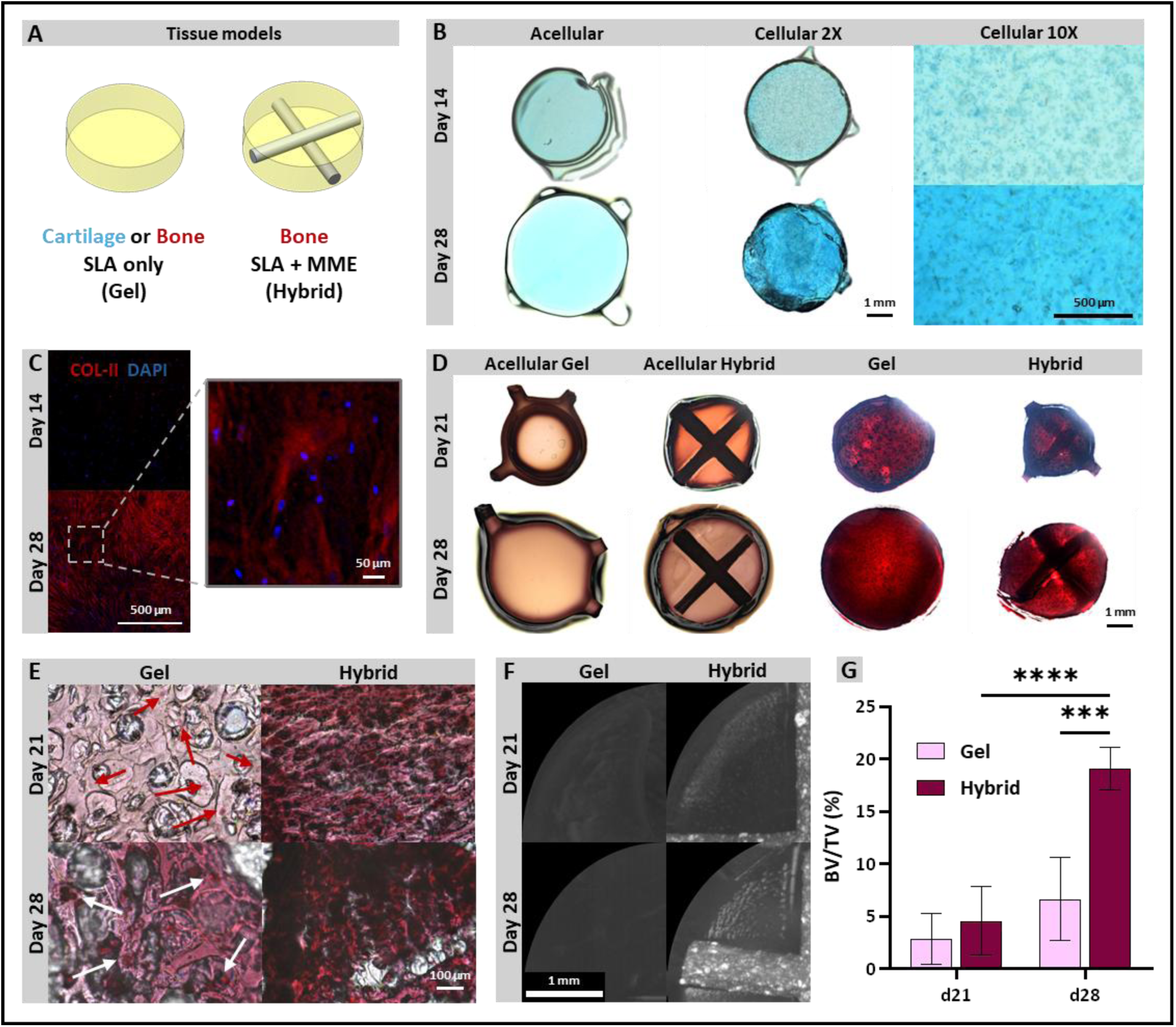
Osteogenic and chondrogenic differentiation of gel and hybrid constructs, composed of GelMA and PCL/β-TCP. (A) Schematics of the bone and cartilage tissue models adapted to the Hybprinter. (B) Chrondrogenic differentiation study demonstrating alcian blue staining of hMSC-laden gel constructs on Day 14 and 28 of differentiation. (C) Collagen type-II (COL-II) immunostaining of hMSC-laden gel constructs on Day 14 and Day 28. Osteogenic differentiation study demonstrating ARS of hMC-laden gel and hybrid constructs at Day 21 and Day 28 of differentiation at (D) 2x and (E) 40x magnification. Red and white arrows indicate positive ARS staining. (F) Maximum projection of microCT images of hMSC-laden gel and hybrid constructs on Day 21 and Day 28. (G) Bone volume/tissue volume (BV/TV) percentage of gel and hybrid constructs.

Additionally, efforts for immunostaining were attempted but unsuccessful in the osteogenic constructs, most likely due to the high density of calcium interfering with the fluorescent microscopy. To further investigate the osteogenic potential of hybrid bioprinting, micro-CT was performed to compare and quantify calcification in gel and hybrid constructs, which is referred to as bone mineralization^60^. Micro-CT images reveal enhanced bone mineralization in hybrid constructs for Days 21 and 28 (**Figure 6F**). Furthermore, bone volume (BV) was quantified and normalized to tissue volume (%) for gel and hybrid constructs, which showed similar levels of mineralization at day 21 for both groups (**Figure 6G**). By Day 28, however, mineralization in gel constructs did not significantly increase whereas mineralization in hybrid samples increased by nearly fivefold. At 28 days of differentiation, nearly 20% of the hybrid constructs was mineralized in comparison to 7% in gels. Note, the mineralization of the cell-laden hybrid constructs were subtracted by acellular hybrid constructs to disregard any mineralization provided by the PCL/β-TCP struts. Histology and micro-CT imaging support the Hybprinter’s enhanced osteogenic potential, highlighting their translational relevance for bone tissue engineering applications.

Here, we demonstrate that the Hybprinter’s capacity to integrate both soft and hard materials into bioprinted constructs allows for advanced musculoskeletal tissue models that closely resemble the stiffness range found in native tissues. The successful billineage differentiation of hMSCs in hybprinted constructs highlights the Hybprinter’s potential to develop multiple types of tissues which can be further enhanced by introducing bioactive materials, such as PCL/β-TCP. Henceforth, the Hybprinter serves as a innovative tool for engineering multi-tissue constructs with unique biomechanical properties for suturable implants, tissue modeling, and therapeutic technologies.

### 2.6. Limitations

Since the Hybprinter was optimized for hybrid bioprinting involving multiple modules, it exhibits certain limitations compared with conventional single-module bioprinters such as syringe-based microextrusion (SBM) or single DLP-SLA systems. First, the integration of two collaborative modules increases the printer’s overall size, occupying approximately 1 m³ of space - larger than typical setups. Second, in comparison with commonly used SBM systems, biomaterial costs are slightly higher, and some unique SBM capabilities, such as embedded printing, are not currently supported. However, the Hybprinter platform is designed to expand and integrate additional modules, particularly SBM and inkjet, in future work. Moreover, similar to most DLP-SLA systems, vat polymerization requires greater amounts of cells and biomaterials compared to SBM systems, nevertheless, it provides enhanced resolution compared to SBM. Third, as detailed in the Discussion section, the MME module increases the distance between the SLA light source and the build plate relative to conventional SLA bioprinters, a constraint addressed by extending exposure times. Lastly, the MME module currently uses a single nozzle, though the Hybprinter’s hardware and software can support dual- and multi-nozzle printing, offering further potential for multi-material scaffold fabrication in upcoming developments.

### 2.7. Conclusion

In this study, we introduced the Hybprinter platform as a solution to unmet needs in complex tissue engineering and personalized regenerative medicine. The system’s biomechatronics and performance were characterized, demonstrating its ability to simultaneously fabricate hybrid constructs that integrate load-bearing scaffolds with multiple hydrogels. Biocompatible materials - including thermoplastics (PLA, PCL, PCL-TCP) and hydrogels (GelMA, PEGDMA, HAMA) - were implemented with diverse cell types (MSCs, HUVECs) under optimized conditions. The hybrid constructs achieved over a 1000-fold increase in mechanical strength compared with hydrogel-only constructs, with a tunable Young’s moduli spanning from 0.014 MPa to 200 MPa, and uniquely exhibited suturability. Biocompatibility was validated with >80% cell viability and proliferative growth over 14 days, while computational and experimental analyses confirmed negligible thermal effects during MME printing. Representative applications included microfluidics, limb-on-a-chip, joint models, vascularized bone grafts, and vertebra-disc replacements. The constructs further demonstrated osteogenic and chondrogenic bilineage differentiation, supporting their musculoskeletal relevance. Collectively, the Hybprinter enables hybrid constructs that reconcile essential attributes of tissue engineering - mechanical robustness, suturability, multi-material integration, gradient properties, and bioactive delivery - highlighting its potential for organ-on-demand applications with patient specificity. While this study illustrates several applications, the platform’s potential extends beyond these demonstrations, with future work directed toward complex multi-tissue models for biological discoveries, therapeutic implants for clinical treaments, and next-generation hybrid bioprinting.

### 2.8. Experimental Section/Methods

*Hybprinter mechatronic design and construction:* The Hybprinter is designed and constructed in-house, incorporating hardware, software, and firmware components. The hardware system consists of a robust base frame, the MME module, a DLP-SLA module, a multi-hydrogel handling system, a motion and positioning system, a temperature control system, microcontroller boards, various electronic components, a power supply, and a computer. The Hybprinter base frame is constructed from slotted aluminum profiles, utilizing a gantry system with Cartesian coordinates for positioning. The positioning system features a timing belt-driven mechanism for XY movement of the MME module, alongside a lead screw-driven system for Z-axis movement of the build plate. Linear motion bearings provide precise guidance, while end stops with limit switches on the X, Y, and Z axes establish operational boundaries and facilitate homing calibration. Motion in all directions is actuated by NEMA17 stepper motors, controlled via G-code from the master program. The electronic systems are powered by a 575W power supply operating at 5V and 12V.

The Hybprinter and its bioprinting process are controlled by a custom-developed MATLAB program on a Windows laptop. This program manages the bioprinting operations, controlling the MME module and multi-hydrogel handling system via G-code transmitted through USB to RAMPS 1.4 microcontroller boards, while also operating the SLA module by projecting photomasks through a projector connected via HDMI. Marlin firmware is employed on the microcontrollers. Pre-print model preparation begins with CAD modeling of the scaffold and hydrogel components in SolidWorks or Autodesk Fusion, with models exported as STL files. For the MME module, the CAD model is processed through Repetier-Host to generate G-code. For the SLA module, Slic3r used to slice STL files into vector graphic SVG files, which are then converted into PNG photomasks using GIMP for projector compatibility. Additionally, a customized MATLAB program is developed to automate the slicing and file conversion processes.

The MME module comprises a hot end nozzle, heating block, heat break, temperature sensor, feeder gear, extruder stepper motor, cooling fan, and a heat sink. The extruder feeds the filament into the hot end, where it is melted and subsequently extruded through the nozzle onto the build plate. The MME module utilizes a 400 μm nozzle to extrude molten thermoplastic filaments with a 1.75 mm diameter onto the build plate following a controlled predefined path to create a porous mesh structure. The printing temperatures of 130°C, 160°C, and 200°C are used for PCL, PCL/β-TCP, and PLA, respectively. When printing scaffolds using the MME module, varying levels of porosity are established during the CAD slicing process, ranging from 15% to 100% based on the specific application. To prevent damage to bioactive components, the build plate is not heated; instead, print bed adhesion is enhanced by reducing the printing speed.

The SLA module features a customized 4K visible-light DLP projector (Optoma 4K UHD) with a brightness of 3000 ANSI lumens and a contrast ratio of 1,000,000:1. The throw distance of the projector is customized and optimized to 20 to 50 cm by increasing the distance between the DLP unit and the projection lens using a spacer. This throw distance refers to the distance from the projector lens to the surface (i.e., build plate) where the image is projected in full focus. Each hydrogel layer is fabricated by projecting a photomask for a predetermined duration, known as the exposure time, onto the regions designated for hydrogel formation. The photomask, or digital mask, is a 2D image where hydrogel regions are represented by white pixels against a black background. The illuminance of the projected light on the Hybprinter build plate is measured at 184,000 Lux. To print the hydrogel, the SLA module lowers the build plate into the vat of liquid polymer, then raises it until a thin layer of liquid remains above the print area, after which the photomask is projected with visible light to photocrosslink the bioink. Before and after each print, the build plate and vats are sterilized with ethanol and then exposed to UV light for 5 minutes, while the printing chamber is also sterilized with ethanol.

The multi-hydrogel handling system features a rotary platform that facilitates the printing of multiple materials. The SLA module operates by lifting the build plate out of one material vat, rotating the rotary table to position another vat beneath the build plate, and then lowering the build plate into the new material. The liquid polymers are contained in biocompatible disposable containers secured to the rotary turntable. The system can autonomously switch between 6 to 12 uniformly spaced vats (ranging from 25 mL to 95 mL each) containing different bioinks for SLA printing, with one or more vats designated for washing during bioink transitions. A robust bearing system maintains the turntable’s level, while a stepper motor facilitates rotation, and a hard stop with a limit switch defines boundaries and calibrates the home position. This design enhances the versatility and efficiency of multi-hydrogel printing. The Hybprinter, along with its associated printing parameters, allows for the adjustment of MME layer thickness between 300 to 600 µm and SLA layer thickness ranging from 100 to 300 µm. The system can implement varying ratios of hydrogel to scaffold layer thickness, ranging from 1:1 to 1:3, which directly correlates to the number of MME and SLA layers, as discussed previously^28^.The layer thickness for both the MME and SLA modules is determined during the pre-print slicing process and G-code generation. SLA layer thickness can be controlled by adjusting exposure time, and the concentration of hydrogel, photo initiator, and photo absorber. In this study, exposure time is primarily utilized to regulate photocrosslinking and SLA layer thickness, while other parameters are optimized to ensure the biocompatibility and non-toxicity of the hydrogel, as detailed in the Supplementary materials. The ratio of MME to SLA layers can be tailored to specific applications, requiring further tuning to meet the desired outcomes.

#### Resolution study

The resolution of the Hybprinter SLA module was evaluated by printing line patterns using a non-diffusible, green fluorescent dye incorporated into PEGDMA pre-polymer solution. A series of linear patterns with decreasing line widths (n=3) were printed and examined under a fluorescent microscope. The smallest successfully fabricated line width, defined as the resolution, below which no lines could be printed.

#### Materials Used

For the MME module of the Hybprinter, the following hard materials were used: 1.75mm PLA filament (Overture 3D Technologies LLC, Houston, Texas), 1.75mm PCL filament (3D4Makers B.V., Haarlem, The Netherlands), and PCL/β-TCP. PCL/β-TCP was engineered at a ratio of 80/20 (PCL to TCP) using in-house methods^34^. Briefly, medical-grade polycaprolactone (Sigma-Aldrich) and β-TCP nanopowder (Berkeley Advanced Materials, Inc.) were weighed with a ratio of 80:20. The weighed PCL pellets were dissolved in DMF overnight at 37°C. The following day, weighed β-TCP powder was added to the solution and left mixing for another hour. The PCL/β-TCP solution was precipitated in 4 L of water and thoroughly rinsed. The precipitated scaffold was then air dried for 24 h at room temperature. The PCL/β-TCP scaffolds were then manually cut into <5 mm pellets and then extruded into filaments using a filament extruder with a 1.75mm diameter nozzle, stainless-steel barrel, and custom-sized screw (Noztek, West Sussex, UK). The extruded filament was then wound into a spool and dried in a seal container prior to 3D bioprinting.

For the SLA module of the Hybprinter, the following soft materials were used: GelMA, PEGDMA (Polysciences 15178-100), and HAMA (Advanced Biomatrix #5212). GelMA was synthesized as previously described ^61^. Briefly, 10% w/v gelatin from porcine skin (Sigma-Aldrich G2500) was dissolved in deionized water at 65°C and stirred. Methacrylic anhydride (Sigma-Aldrich 276685) was added dropwise to the solution at a molar ratio of 100:1 (methacrylic anhydride: gelatin) and stirred for five hours, while protected from light. Dialysis was performed for five days against DI water using a dialysis tube (Spectrum Laboratories, Rancho Dominquez, CA), then the resulting aqueous solution was vacuum filtered and lyophilized for three days and stored at –80°C. Methacrylation of synthesized GelMA was confirmed with a ^1^H-NMR spectra performed with an Bruker Avance Neo 500 MHz NMR spectrometer^62,63^. For all non-cellular constructs, PEGDMA was prepared at 30% w/v in DI water and/or PLA filament was prepared.

#### Mechanical Characterization

All mechanical tests were performed using an Instron 5944 uniaxial testing system (Instron Corporation, Norwood, MA). Depending on the loading range, appropriate load cells with capacities of 2 N, 10N, 100 N, and 2 kN were employed, with a preload ranging from 1 mN to 0.1 N and a displacement rate of 0.1 mm/s, up to a maximum strain of 40%. Stiffness was calculated from a linear curve fit applied to the near-linear region of the stress-strain curve within the 0%–20% strain range. The tensile and compressive moduli of PEGDA hydrogel samples fabricated with the Hybprinter were determined through compression tests on disks with a diameter of 12 mm and a height of 4 mm, and tensile tests on dog-bone shaped samples, with three samples per group. The elastic modulus of GelMA hydrogel samples fabricated by Hybprinter at varying exposure times was determined through compressive tests. Disks with a diameter of 8 mm and a height of 3 mm were printed using exposure times of 60, 90, 120, and 150 seconds. Four samples per group underwent compression testing. The Young’s modulus of the three thermoplastic materials used in the Hybprinter MMA module – PLA, PCL, and PCL/β-TCP – was assessed through tensile testing on filament specimens with 50 mm in length and 1.75 mm in diameter, with four samples per group The mechanical properties of hybrid samples printed with the Hybprinter were assessed through compression and tensile testing. For compression testing of hybrid samples, square samples measuring 10 × 10 mm with a height of 4.5 mm and 85% porosity were fabricated. For tensile testing of hybrid samples, rectangular samples measuring 13 × 10 mm with a height of 1.5 mm and 85% porosity were printed. The hybrid samples were fabricated using PLA as the scaffold and PEGDA as the hydrogel (n=3).

#### Suture Retention Test

The suture retention test was conducted on hybrid samples consisting of a rectangular scaffold measuring 30 × 10 mm with a height of 1.5 mm and 85% porosity. The central 10 × 12 mm region of the scaffold was integrated with hydrogel, while the two ends were left for handling during the suturing process. Three suture ties were applied to each end of the sample using Ultrabraid size 2 suture thread. The test was performed under tensile force until failure. A similar suture retention test was conducted on scaffolds without hydrogel for comparison (n=3).

#### Heat Transfer Computational Analysis

The effect of 3D printing using heated thermoplastic material on hydrogel is studied by creating a computational heat transfer (CHT) model and simulation in COMSOL (COMSOL Inc. USA). The CHT model is a finite element (FE) model representing the 2D cross-section of the hybrid bioprinting environment. It incorporates hydrogel and air sections at the bottom and top of the solution domain, respectively, along with a circular section representing the printed hot thermoplastic strut with a diameter of 400 μm, with the initial condition set at the initial extrusion temperature from the nozzle. Owing to different printing temperatures for different materials, three separate models associated with the thermoplastic materials employed by the Hybprinter, namely, PLA, PCL, and PCL-TCP, were configured and simulated, corresponding to printing temperatures of 200°C, 130°C, and 160°C, respectively. The hydrogel thermal properties were approximated to those of water. The boundaries of the model are assumed to be at a constant room temperature of 22°C. The entire solution domain mesh was defined as a standard, physics-controlled mesh with triangular elements. The time-dependent dynamic solution of the CHT simulation was obtained using the finite element method (FEM). The variations in hydrogel temperatures at two distinct distances (100 and 200 μm) from the strut at three distinct points were obtained and compared across the three material-dependent models. The simulation results illustrate the temperature distribution within the hydrogel and scaffold sections, the maximum temperature at each measurement point, and the temporal temperature variation.

#### Cell culture

Bone marrow-derived human mesenchymal stem cells (hMSCs, Lonza PT-2501) were used for biocompatibility studies. Cells were cultured in mesenchymal stem cell basal medium supplemented with mesenchymal cell growth serum, L-glutamine, and gentamicin following manufacturer’s instructions (Lonza PT-3001). Cells were incubated at 37°C and 5% CO2. The medium was replenished every 3 – 4 days, and cells were passaged at 90% confluency. For exposure time experiments, cells were used on passages < 18. For all other experiments, cells were used on or before passage 8. Human umbilical vascular endothelial cells tagged with green fluorescent protein (gHUVECs) and blue fluorescent protein (bHUVECs) were gifted by Prof. Roger Kamm at the Massachusetts Institute of Technology. Cells were cultured in VascuLife VEGF Endothelial Medium (Lifeline Cell Technology LL-0003), along with the manufacturer’s recommended growth factors. Cells were incubated 37°C and 5% CO2. The medium was replenished every 2 – 3 days, and cells were passaged at 80 – 90% confluency.

#### Preparation of bioink for cellular constructs

Lyophilized GelMA was reconstituted at 10% w/v in PBS (Gibco 10010-023) and placed in a shaking incubator at 37°C. The GelMA solution was then centrifuged at 3000 RCF for 10 minutes to remove any foam. Lithium phenyl-2,4,6-trimethylbenzoylphosphinate (LAP) (Sigma Aldrich 900889) was added as a photoinitiator at a final concentration of 0.3% and tartrazine (Sigma Aldrich T0388) was added as a photoabsorber at a final concentration of 0.02%^29,64^. The GelMA bioink was placed in a shaking incubator at 37°C for 15 minutes after adding each reagent, and an additional centrifugation step was performed if foam was present in the solution. For cellular studies, an additional sterile filtering step was performed with a 0.22 µM PVDF syringe filter, prior to adding cells. Before loading cells into bioink, cells were detached from the flask using Trypsin/EDTA. Once cells were detached, cell culture medium was added in a 6:1 ratio (medium: trypsin) and centrifuged at 500 x g for 5 minutes at room temperature. Lastly, the supernatant was removed, and the remaining pellet was homogenized with GelMA. All hMSC-laden bioinks were prepared with a final cell density of 1 x 10^6^ cells mL^-1^ unless stated otherwise.

#### Viability and proliferation assays

To assess the biocompatibility of the Hybprinter, viability and proliferation assays were performed on cellular constructs every 48 hours for one week. Samples with 6mm diameter and 1.5 mm of thickness were printed (n=4). Immediately after printing, cellular constructs were transferred to a 24 well plate (one per well). Inside a biosafety cabinet, cellular constructs were washed with PBS three times prior to adding 500 µL of cell culture medium and incubating at 37°C and 5% CO2. For viability, live cells were stained with calcein AM and dead cells were stained with ethidium homodimer-1 (Invitrogen L3224), following manufacturer’s instructions. After staining, cellular constructs were protected from light and incubated at 37°C for 45 minutes. Fluorescent images were then taken with a Keyence BZ-X810 microscope. Viability was quantified by counting the number of live and dead cells and viability ratio was calculated with live cells: total number of cells. For viability assays in hybrid constructs, a two-way Analysis of Variance (ANOVA) was performed to evaluate statistical significance of viability between the core and perimeter regions and its timepoints. For all other viability assays, a one-way ANOVA was performed instead. For proliferation studies, an MTS assay was performed following manufacturer’s instructions (Abcam ab197010). Once the MTS reagent was added, cellular constructs were protected from light and incubated at 37°C for two hours. Then, cell proliferation was quantified via absorbance using a plate reader (SpectraMax iD3). Numerical data was normalized against acellular constructs.

#### Multi-cellular and multi-material bioprints

Multi-cellular constructs were bioprinted using a final cell density of 0.7 x 10^6^ cells mL^-1^ for all fluorescent markers. Cells were printed in 10% GelMA while surrounding acellular gel was printed in 10% PEGDMA. Fluorescent images and Z-stacks of the constructs were taken with a Keyence BZ-X810 immediately following the printing process. Images were post-processed with ImageJ.

#### Microchannel printing

For microchannel printing, a 1% HAMA solution was prepared in PBS without calcium or magnesium. Channels of 1 mm width were printed with HAMA at an exposure time of 100 s via SLA. For enzymatic degradation of HAMA, a 150 U/mL of hyaluronidase (STEMCELL Technologies #07461) was prepared in PBS and sterile filtered. The constructs were submerged in the enzymatic solution for 72 hr at 37°C prior to flushing out the degraded gel.

#### Chondrogenic differentiation

hMSC-laden gels were bioprinted at a final density of 2.5 x 10^6^ cells ml^-1^ in 10% GelMA via SLA. Acellular gels were printed on the same day following the same conditions. All constructs were post-washed with PBS three times after printing. Cellular constructs were incubated at 37°C in chondrogenic differentiation medium (A1007101) and underwent medium changes every 2 – 3 days for 28 days. Acellular constructs were incubated in PBS under the same conditions.

#### Osteogenic differentiation

hMSC-laden gels and hybrid constructs were bioprinted at a final density of 1.0 x 10^6^ cells ml^-1^ in 10% GelMA via SLA. For hybrid constructs, PCL/β-TCP filament was printed via MME within the gel. Acellular gels and hybrid constructs were printed on the same day following the same conditions. All constructs were post-washed with PBS three times after printing. Cellular constructs were incubated at 37°C in osteogenic differentiation medium (A1007201) and underwent medium changes every 3 – 4 days for 28 days. Acellular constructs were incubated in PBS under the same conditions.

#### Histology

Alcian Blue staining was performed for chondrogenic constructs on Day 14 and Day 28. Samples were fixed in 4% paraformaldehyde for 45 minutes at room temperature, followed by three PBS washes. Fixed samples were stained with 1% Alcian Blue solution prepared in 0.1 N HCl for 45 minutes. Chondrogenic samples were then rinsed three times with 0.1 N HCl, followed by multiple washes of DI water (5 – 10 minutes each) until the solution appeared mostly clear. Chondrogenic samples were left overnight shaking moderately at 4°C in 3% acetic acid. The following day, samples were then rinsed three times with DI water prior to imaging. Alizarin Red Staining was performed for osteogenic constructs on Day 21 and Day 28. Samples were fixed in 4% paraformaldehyde for 45 minutes at room temperature, followed by three PBS washes. Samples were incubated with 2% Alizarin Red Stain Solution (CM-0058) for 30 minutes at room temperature then washed gently with DI water until solution turned mostly clear. Samples were left gently shaking in DI water for 72 hours at 4°C prior to imaging. For slicing, samples were submerged in Optimal Cutting Temperature Embedding Medium (Fisher 4585) and stored overnight at -80°C. 100-µm slices were extracted from the samples and mounted onto a microscope glass slide using a Leica CM3050 S cryostat set at -20°C. All histology images were taken with a Keyence BZ-X810. All images were postprocessed with ImageJ.

#### Immunofluorescence

Multiple overnight steps were incorporated to ensure proper diffusion of antibodies and fluorescent dyes within the 3D constructs. Chondrogenic and osteogenic samples were collected and fixed in 4% paraformaldehyde, washed three times with PBS, then temporarily stored in 30% sucrose in PBS at 4°C and protected from light until staining. Fixed samples were permeabilized and blocked with 0.5% Triton X-100 in 3% bovine serum albumin (BSA) (sc-2323), followed by three PBS washes. Primary antibodies were diluted 1:200 and 1:100 in 0.1% Triton X-100 and 3% BSA for collagen type II (ab34712) and osteocalcin (sc-365797), respectively. Samples were incubated in their respective primary antibodies and left shaking gently overnight at 4°C, followed by three PBS washes. The next day, secondary antibodies were diluted 1:500 in 0.1% Triton X-100 and 3% BSA for Alexa Fluor 647 (711-605-152) and Alexa Fluor 488 (ab150113). Samples were incubated in their respective secondary antibodies and left shaking gently overnight at 4°C, followed by three PBS washes. The next day, samples were incubated with 2 µg/mL of DAPI and left shaking gently overnight at 4°C. To remove any background noise, samples were washed three times with 0.1% Tween-20 in PBS and then incubated with 0.05% Tween-20 in PBS and left shaking gently overnight at 4°C. On the final day, samples were mounted on a microscope glass slide and images were taken with a Leica STELLARIS 5 confocal microscope. All images were postprocessed in ImageJ.

#### MicroCT

Calcium deposits in osteogenic constructs were evaluated at Day 21 and Day 28 by micro computed tomography (microCT). Samples were collected and fixed in 4% paraformaldehyde, washed three times with PBS, then temporarily stored in PBS at 4°C until imaging. Samples were loaded onto the microCT platform along with a rat cancellous bone phantom for reference. Images were taken with a Bruker SkyScan 1276 with a pixel size of 3.99 µm, 360° rotation, and a rotation step of 0.150°. MicroCT images were reconstructed and analyzed using the Bruker SkyScan suite and Z-stacks were generated with ImageJ. For bone volume quantification, the bone volumes for cellular hybrid samples were subtracted against the bone volumes of acellular hybrid samples to disregard bone mineralization from the PCL/β-TCP filaments. Then, bone volume was normalized to hybrid samples’ tissue volume. A two-way ANOVA was performed to evaluate statistical significance of bone volume between hybrid/gel constructs and timepoints.

#### Statistical analysis

Reported values for experimental outcomes are expressed as mean ± standard deviation. Statistical evaluations employed a one-way ANOVA, followed by a post-hoc Tukey’s test, unless stated otherwise. A threshold of p < 0.05 was utilized to determine statistical significance.

**Figure S1.**
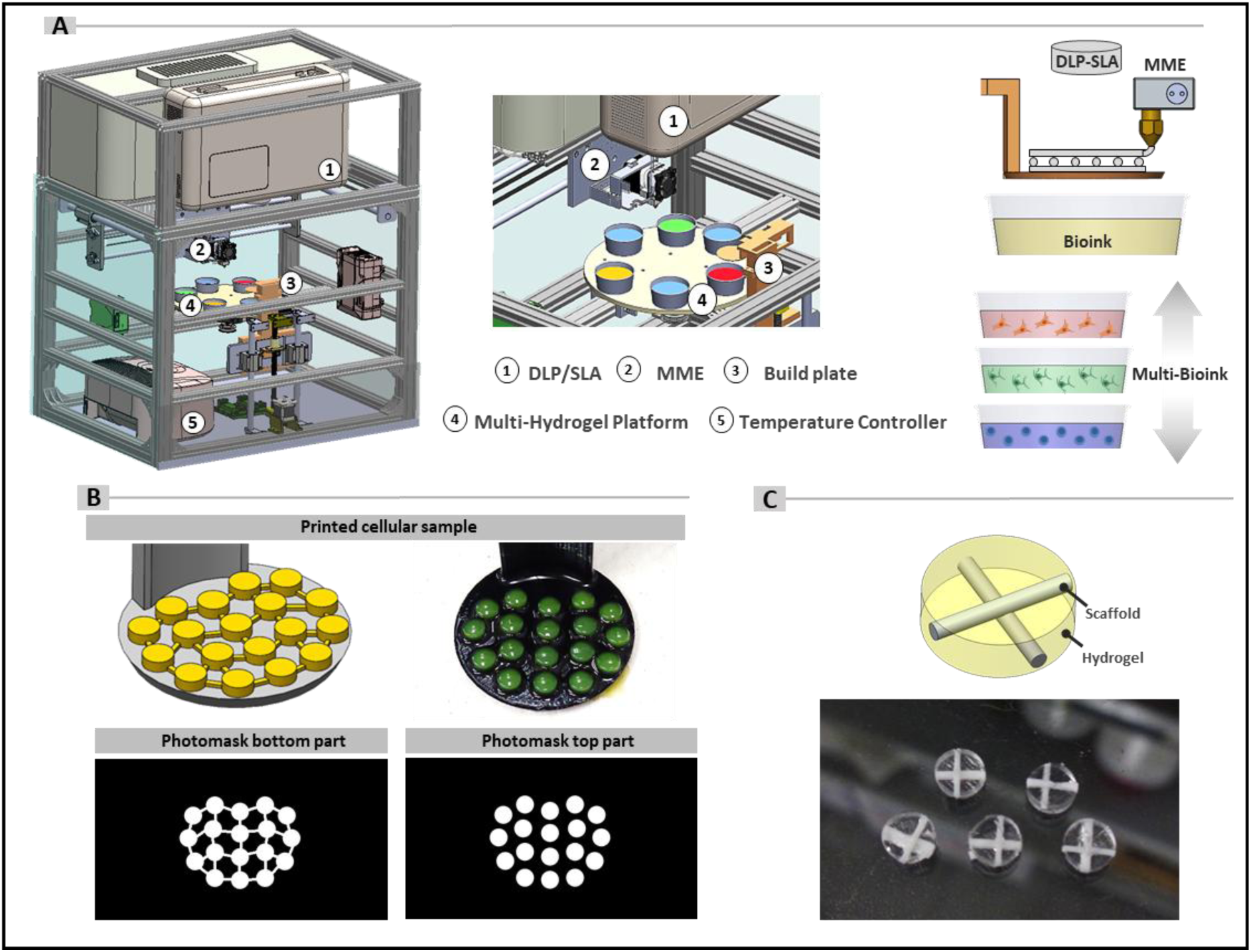
Hybprinter and representative prints. (A) Overview of the Hybprinter design and its main components. (B) Printed MSC-laden GelMA disks with associated photomasks, illustrating the link structure in the first two SLA layers to enhance stability and adherence to the build plate. (C) Representative hybrid disks fabricated using the Hybprinter.

**Figure S2.**
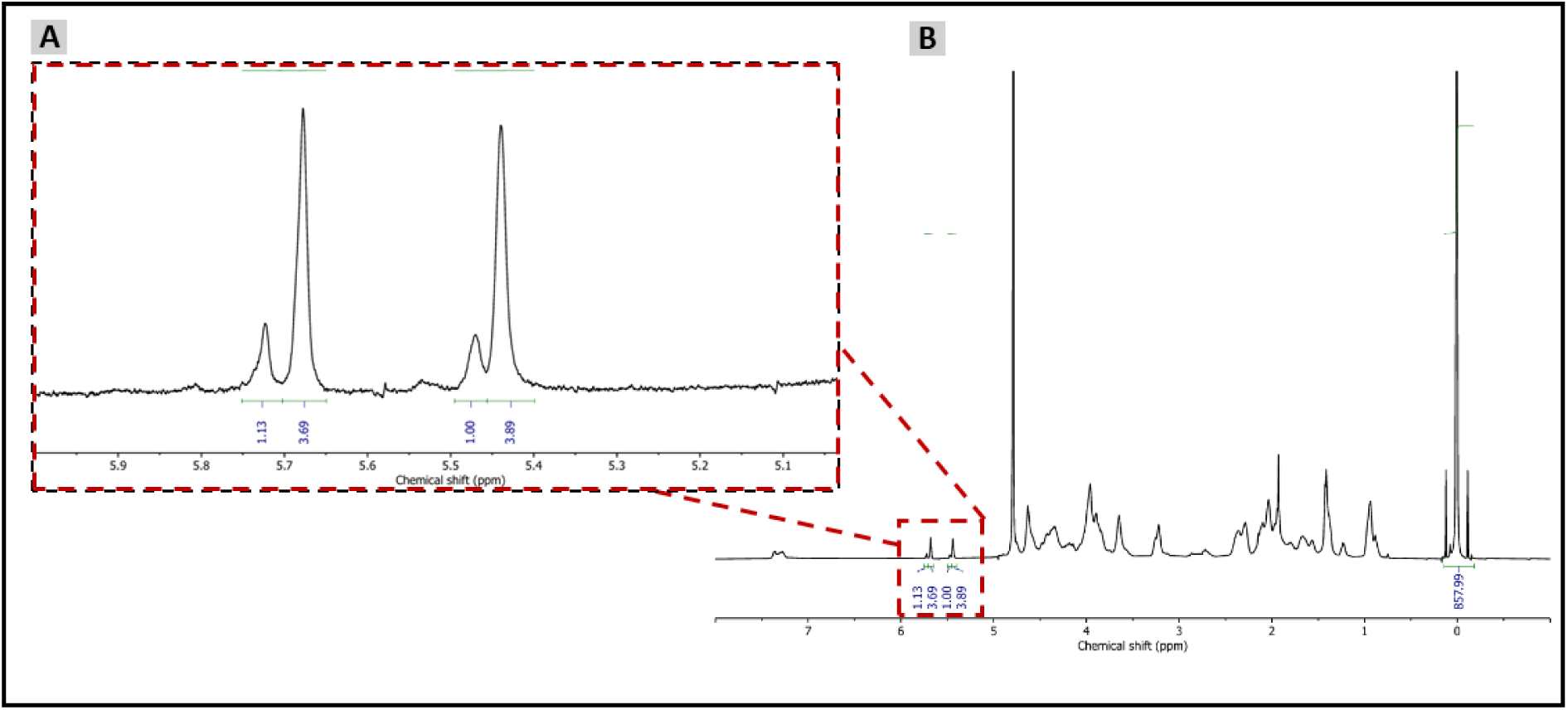
^1^H-NMR spectra of synthesized GelMA. **A)** Acrylic protons of methacrylamide groups in lysine and hydroxylsine residues in GelMA are highlighted between 5 and 6 ppm. TMSP as a reference point is highlighted at 0ppm. **B)** Zoomed in ^1^H-NMR spectra of GelMA from 5.1 to 5.9 ppm.

**Figure S3.**
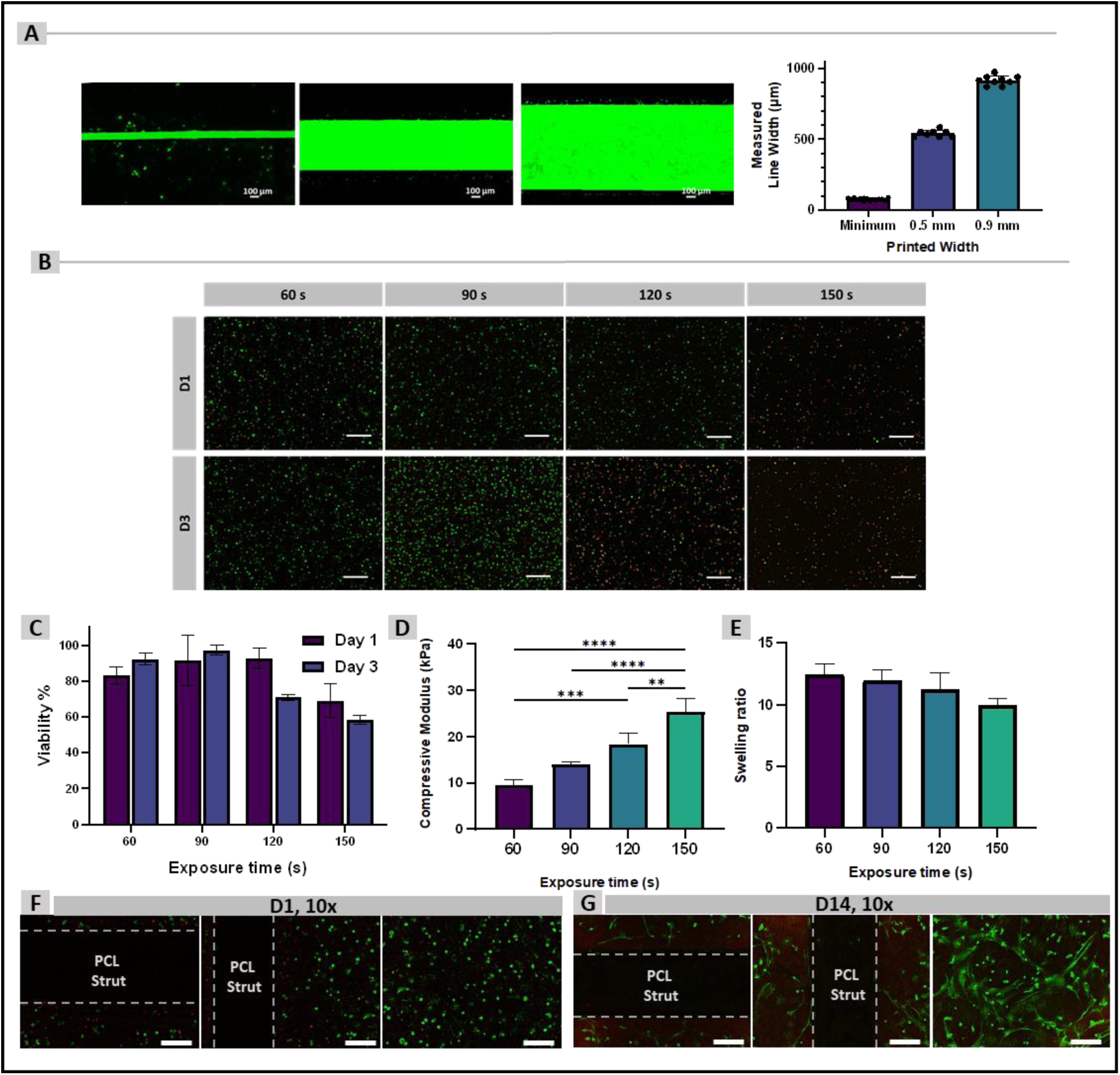
Characterization of the Hybprinter SLA module and magnified images of hybrid samples. (A) Resolution assessment prints determining the minimum line width achievable by the Hybprinter SLA module. (B) Live-dead fluorescent images of MSC-laden GelMA samples at different exposure times for optimization of the SLA printing process. (C) Viability ratio at various SLA exposure times. (D) GelMA compressive modulus measured at different exposure times. (E) GelMA swelling ratio at different exposure times. (F) Live-dead fluorescent images of the perimeter and core regions of hybrid samples on Day 1. (G) Live-dead fluorescent images of the perimeter and core regions of hybrid samples on Day 14.

## Acknowledgements

We gratefully acknowledge all funding support provided to Professor Yunzhi Peter Yang, including R01AR072613 and R01AR074458 (NIAMS), and R01CA268514 (NCI). We also would like to thank the National Science Foundation’s Graduate Research Fellowship Program and Stanford Bio-X’s Fellowship for their sponsorship and funding to ASFP. Special thanks to Bruce and Elizabeth Dunlevie for their generous donation and support as part of the Stanford Bio-X Fellowship program. We would also like to acknowledge The Nucleus facility at Stanford University for their training and support in NMR studies (Bruker Avance Neo 500 MHz NMR spectrometer NIH High End Instrumentation grant (1 S10 OD028697-01)). We would like to thank Mr. Calvin Chan from the Department of Orthopaedic Surgery at Stanford University for his assistance in mechanical testing. We would also like to acknowledge the Stanford Center for Innovation in *In vivo* Imaging (SCi^3^) small animal imaging center for assistance and technological support in microCT imaging and data analysis (NIH S10 Shared Instrumentation Grant 1S10OD02349701). Special thanks to Prof. Roger Kamm and Dr. Zhengpeng (Jason) Wan from the Kamm research group at the Massachussetts Institute of Technology for their support and generous donation of fluorescently-tagged HUVECs.

